# Neuroanatomical Changes in White and Grey Matter after Sleeve Gastrectomy

**DOI:** 10.1101/678284

**Authors:** Andréanne Michaud, Mahsa Dadar, Mélissa Pelletier, Yashar Zeighami, Isabel Garcia-Garcia, Yvonne Yau, Mélanie Nadeau, Simon Marceau, Laurent Biertho, André Tchernof, D. Louis Collins, Denis Richard, Alain Dagher, for the REMISSION Study Group

## Abstract

**Background:** MRI studies show that obese adults have reduced grey (GM) and white matter (WM) tissue density as well as altered WM integrity. It remains to be examined if bariatric surgery induces structural brain changes. The aim of this study is to characterize GM and WM density changes in a longitudinal setting, comparing pre- and post-operation and to determine whether these changes are related to inflammation and cardiometabolic markers.

**Methods:** 29 severely obese participants (age: 45.9±7.8 years) scheduled to undergo sleeve gastrectomy (SG) were recruited. High-resolution T1-weighted anatomical images were acquired 1 month prior to as well as 4 and 12 months after surgery. GM and WM densities were quantified using voxel-based morphometry (VBM). Circulating lipid profile, glucose, insulin and inflammatory markers (interleukin (IL)-6, C-reactive protein (CRP) and lipopolysaccharide-binding protein (LBP) were measured at each time point. A linear mixed effect model was used to compare brain changes before and after SG, controlling for age, gender, initial BMI and diabetic status. To assess the associations between changes in adiposity, metabolism and inflammation and changes in GM or WM density, the mean GM and WM densities were extracted across all the participants using atlas, and linear mixed-effect models were used.

**Results:** As expected, weight, BMI, waist circumference and neck circumference significantly decreased after SG compared with baseline (p<0.001 for all). A widespread increase in WM density was observed after surgery, particularly in the cerebellum, brain stem, cerebellar peduncle, cingulum, corpus callosum and corona radiata (p<0.05, after FDR correction). Significant increases in GM density were observed 4 months after SG compared to baseline in several brain regions such as the bilateral occipital cortex, temporal cortex, precentral gyrus and cerebellum as well as right fusiform gyrus, right hippocampus and right insula. These GM and WM increases were more pronounced and widespread after 12 months and were significantly associated with post-operative weight loss and the improvement of metabolic alterations. Our linear mixed-effect models also showed strong associations between post-operative reductions in LBP, a marker of inflammation, and increased WM density. To confirm our results, we tested whether the peak of each significant region showed BMI-related differences in an independent dataset (Human Connectome Project). We matched a group of severely obese individuals with a group of lean individuals for age, gender and ethnicity. Severe obesity was associated with reduced WM density in the brain stem and cerebellar peduncle as well as reduced GM density in cerebellum, regions that significantly changed after surgery (p<0.01 for all clusters).

**Conclusions:** Bariatric surgery-induced weight loss and improvement in metabolic alterations is associated with widespread increases in WM and GM densities. These post-operative changes overlapped with baseline brain differences between severely obese and normal-weight individuals, which may suggest a recovery of WM and GM alterations after bariatric surgery.

## INTRODUCTION

Adding to the well-known metabolic alterations related to obesity such as insulin resistance, dyslipidemia, elevated blood pressure and chronic low-grade inflammation (1, 2), epidemiological studies and recent meta-analyses reported that moderate lifetime overweight/obesity is related to an increased risk of cognitive impairment and incident dementia (3–5). Brain alterations induced by obesity may underlie the link between obesity and cognitive dysfunction (6, 7). Thus, a better understanding of the structural brain changes associated with obesity and body weight fluctuation in humans is of particular relevance.

MRI studies show that obese adults have reduced grey matter (GM) and white matter (WM) tissue density. For instance, our recent neuroimaging coordinate-based meta-analysis on voxel-based morphometry (VBM) demonstrated that obesity is associated with reduced GM volume in discrete brain regions, namely the ventromedial prefrontal cortex, cerebellum and portions of temporal and parietal lobes (8). A study in morbidly obese participants undergoing bariatric surgery found widespread cortical and subcortical GM atrophy prior to surgery (9). In a cross-sectional study, obesity was also associated with a greater degree of WM atrophy, with an estimated increase in brain age of 10 years (10). There is a possible link between obesity-related GM atrophy and dementia, as the BMI-GM atrophy correlation is also seen in individuals with mild cognitive impairment and Alzheimer’s Disease (11).

Findings on WM structure and integrity in obesity are less clear (12, 13). Some studies report a positive association between obesity and WM density in frontal, temporal and parietal regions (6, 14), while other studies find a negative association in basal ganglia and corona radiata (15). WM hyperintensities, which are markers of central nervous system ischemic disease, have also been observed in greater numbers in obese individuals (12, 13, 16). Studies using diffusion tensor imaging have shown a loss of WM integrity linked to cognitive impairments in obese participants (7, 17–19).

The mechanism linking obesity with reductions in brain tissue remains largely unknown. Animal studies suggest that chronic inflammation associated with obesity can lead to adverse effects on brain tissue (20, 21). Intriguingly, recent discoveries in humans point to the chronic low-grade inflammation as a potential mechanism to explain obesity-induced WM alterations (13, 16, 19, 21, 22). However, it is unclear if these brain changes are permanent or whether they can be reversed after weight loss.

Bariatric surgery represents an interesting approach to address these questions, since it allows the examination of the effects of sustained weight loss and long-term metabolic improvements on the brain in a longitudinal setting (23–25). Growing evidence shows improvement in cognitive functions after bariatric surgery (26, 27). However, only few structural MRI studies examined changes in GM and WM densities after bariatric surgery (9, 28, 29). Tuulari et al. (9) were the first to show global increase in WM density and limited increase in GM density (occipital and temporal regions) 6 months after bariatric surgery using VBM. Zhang et al. (29) observed increased GM and WM densities as well as improved WM integrity in regions involved in food intake control and cognitive-emotion regulation 1 month following sleeve gastrectomy (SG) compared to baseline. A recent study found widespread WM and GM density changes in all cortical and subcortical structures as well as the brainstem and cerebellum 1 year after gastric bypass surgery (28). The improvement in metabolic alterations following bariatric surgery-induced weight loss has been suggested as a potential mechanism to explain these brain changes. However, a better understanding of the link between cardiometabolic variables, inflammatory markers and brain structural changes is needed.

In the present longitudinal study, we aimed to characterize GM and WM density changes 4 months and 12 months after SG, comparing pre- and post-surgery and to determine whether these changes are associated with inflammation and cardiometabolic markers. We tested the hypothesis that bariatric surgery and the associated weight loss induce recovery of GM and WM density, and that these changes are related to improvements in inflammatory and metabolic variables.

## MATERIALS AND METHODS

### Participant recruitment

This study was part of the REMISSION trial (Reaching Enduring Metabolic Improvements by Selecting Surgical Interventions in Obese iNdividuals, NCT02390973). The study sample included 29 participants with severe obesity (8 men, 21 women; mean age=45.9±7.8 years; mean BMI=44.1±4.6 kg/m^2^) scheduled to undergo SG at the *Institut universitaire de cardiologie et de pneumologie de Québec*. The indications for surgery followed NIH guidelines (30). Twenty-seven participants completed the experiments both pre-surgery and 4 months after SG. Among those participants, we tested 17 participants 12 months after surgery. Exclusion criteria were the following: 1) BMI<35 kg/m^2^; 2) age under 18 or over 60 years; 3) any uncontrolled medical, surgical, neurological or psychiatric condition; 4) cirrhosis or albumin deficiency; 5) any medication that may affect the central nervous system; 6) pregnancy; 7) substance or alcohol abuse; 8) previous gastric, oesophageal, brain or bariatric surgery; 9) gastro-intestinal inflammatory diseases or gastro-intestinal ulcers; 10) severe food allergy; and 11) contraindications to MRI (implanted medical device, metal fragment in body, or claustrophobia). The Research Ethics Committee of the *Centre de recherche de l’Institut universitaire de cardiologie et de pneumologie de Québec* approved the study. All participants provided written, informed consent to participate in the study.

### Surgical procedures

All patients were operated laparoscopically. A 250 cm^3^ vertical SG starting 5 cm to 7cm from to the pylorus to the Hiss angle was performed using a 34-44 French Bougie for guidance, to create a gastric tube (31). The greater curvature and fundus of the stomach were removed.

### Study design and experimental procedures

Participants were studied approximately 1 month prior to as well as 4 and 12 months after surgery. At each study visit, participants underwent a physical examination including measurement of blood pressure, and fasting blood biochemistry. T1-weighted anatomical images were acquired on the morning (started between 9:00 and 10:30am) of each visit. Participants were asked to fast for 12 hours before the scanning session and received a standardized beverage meal (237ml, Boost original, Nestle Health Science) 1 hour before the MRI session. The standardized beverage contained a total of 240 kcal, 41g of carbohydrates, 10g of proteins and 4g of lipids. It was consumed over a 5-min period. Before each MRI session, body weight and height were measured using standardized procedures to calculate BMI (kg/m^2^) as well as percentage of excess weight loss (%EWL) and percentage of total weight loss (%TWL). The %EWL was calculated using total pre-operative weight, post-operative weight, and ideal body weight (IBW) for an ideal BMI of 23 kg/m^2^ as previously used (32). Waist circumference and neck circumference were also measured at each visit using standardized procedures.

### Plasma lipid profile, glucose homeostasis and inflammatory markers

Blood samples were collected on the morning of each visit after a 12-hour fast in EDTA-coated tubes or serum clot activator tubes. All samples were immediately placed at 4°C and then centrifuged and stored at −80°C. Plasma insulin levels were measured using chemiluminescence immunoassay and plasma glucose levels were measured using the hexokinase method (33). The homeostasis model assessment insulin resistance (HOMA-IR) index was calculated (34). Cholesterol and triglyceride levels in serum and lipoprotein fractions were measured with a Siemens Dimension Vista 1500 using enzymatic methods. Plasma interleukin (IL)-6 and lipopolysaccharide-binding protein (LBP) levels were measured by commercially available enzyme-linked immunosorbent assay (R&D Systems, Minneapolis, MN and HyCult Biotechnology, Huden, the Netherlands, respectively)(22, 35). Plasma CRP concentrations were measured using the high-sensitivity immunonephelometric assay.

### T1-weighted MRI acquisition

T1-weighted three-dimensional (3D) turbo field echo images were acquired at the initial visit, 4 months and 12 months after surgery using a 3T whole-body MRI scanner (Philips, Ingenia, Philips Medical Systems, X) equipped with a 32-channel head coil at the *Centre de recherche de l’Institut universitaire de cardiologie et pneumologie de Québec*. The following parameters were used: 176 sagittal 1.0 mm slices, repetition time/echo time (TR/TE) = 8.1/3.7ms, field of view (FOV)= 240 × 240 mm^2^, voxel size = 1×1×1 mm.

### Voxel-based morphometry measurements

T1-weighted structural scan of each participant was used to measure voxel-based morphometry. GM and WM densities were assessed from each T1-weighted MRI using a standard VBM pipeline (8). The preprocessing steps were the following: 1) image denoising (36); 2) intensity non-uniformity correction (37); and 3) image intensity normalization into range (0-100) using histogram matching. Images were then first linearly (using a nine-parameter rigid registration) and then nonlinearly registered to an average brain template (MNI ICBM152) as part of the ANIMAL software (38) and segmented into GM, WM and cerebrospinal fluid images. These steps remove global differences in the size and the shape of individual brains and transform individual WM or GM density maps to the standardized MNI ICBM152 template space. VBM analysis was performed using MNI MINC tools (http://www.bic.mni.mcgill.ca/ServicesSoftware/MINC) to generate WM and GM density maps representing the local WM/GM concentration per voxel. To avoid the overlap between GM and WM signal in the border of GM and WM (due to a combination of partial volume effect and smoothing of the maps), we remove 3 voxels on the border of the GM and WM regions.

### Statistical analyses

Repeated-measures ANOVA or Student’s paired-t-tests were performed to examine changes in adiposity, metabolic and inflammatory measurements after SG. Linear mixed-effects models were used to assess WM and GM changes following SG, with session (baseline, 4 months or 12 months) as a fixed factor and subject as a random factor. Age, sex, initial BMI, and diabetic status were included in the model as covariates. The mixed-effects model estimates are represented by Beta values in the results section. VBM results were whole-brain False Discovery Rate (FDR) corrected (p<0.05). To examine the associations a adiposity, metabolic and inflammatory variables and surgery-induced brain changes in GM or WM density, the AAL Atlas (39) was used to extract average regional VBM GM values and similarly, the Atlas from Yeh et al. (40) was used to extract regional VBM WM values for each participant. Mean GM and WM densities were extracted across all the participants. Linear mixed-effects models were used to assess the association between changes in adiposity, metabolism and inflammation and changes in GM or WM density (dependent variables). Age and sex were included in the model as covariates. All continuous variables were log-transformed or z-scored prior to the analysis. The Bonferroni correction was applied for multiple comparisons. Linear mixed-effect models were fitted using *fitlme* from MATLAB version R2017a (The MathWorks Inc., Natick, MA). Other statistical analyses were performed with JMP software version 14 (SAS Institute Inc, Cary, NC, USA).

### Validation in an independent dataset from the Human Connectome Project

We used an independent sample from the Human Connectome Project (HCP) to test whether the WM and GM regions that significantly changed after the surgery show BMI-related differences in obese versus normal weight participants in a separate adult population (41). All the participants from the HCP with a BMI higher than 35kg/m^2^ were included in the study. These participants (n=46) were individually matched (1:1) for age, sex and ethnicity with a group of normal weight individuals (n=46) from the HCP. Other exclusion criteria included participants with missing information on age, sex, BMI and ethnicity. T1-weighted 3D MPRAGE sequence with 0.7mm isotropic resolution images were acquired by the HCP investigators using a 3T MRI scanner (Siemens Skyra) equipped with a 32-channel head coil. The following parameters were used: 256 sagittal slices in a single slab, TR/TE = 2400ms/2.14ms, T1=1000 ms, field of view (FOV)= 224 mm, FA=8°, Echo Spacing=7.6 ms, voxel size = 0.7×0.7×0.7 mm^3^ (42). GM and WM VBM were calculated using the same procedure as in our clinical sample. Student’s paired *t*-tests were used to compare peak VBM values of the regions that significantly changed after surgery in our sample between severely obese and normal-weight HCP participants.

## RESULTS

### Clinical characteristics of participants

Clinical characteristics of study participants are shown in **Table 1**. As expected, weight, BMI and waist circumference significantly decreased 4 months and 12 months after SG compared with baseline (p<0.001 for all). The mean %EWL was 46.7±10.3 after 4 months and 68.1±13.4 after 12 months. The mean %TWL was 21.3±4.3 after 4 months and 31.1±5.4 after 12 months. Significant improvements in blood pressure, glucose homeostasis, HDL-cholesterol and triglyceride levels as well as circulating inflammatory markers (CRP, IL-6 and LBP) were observed after surgery (**Table 1**).

**Table 1.**
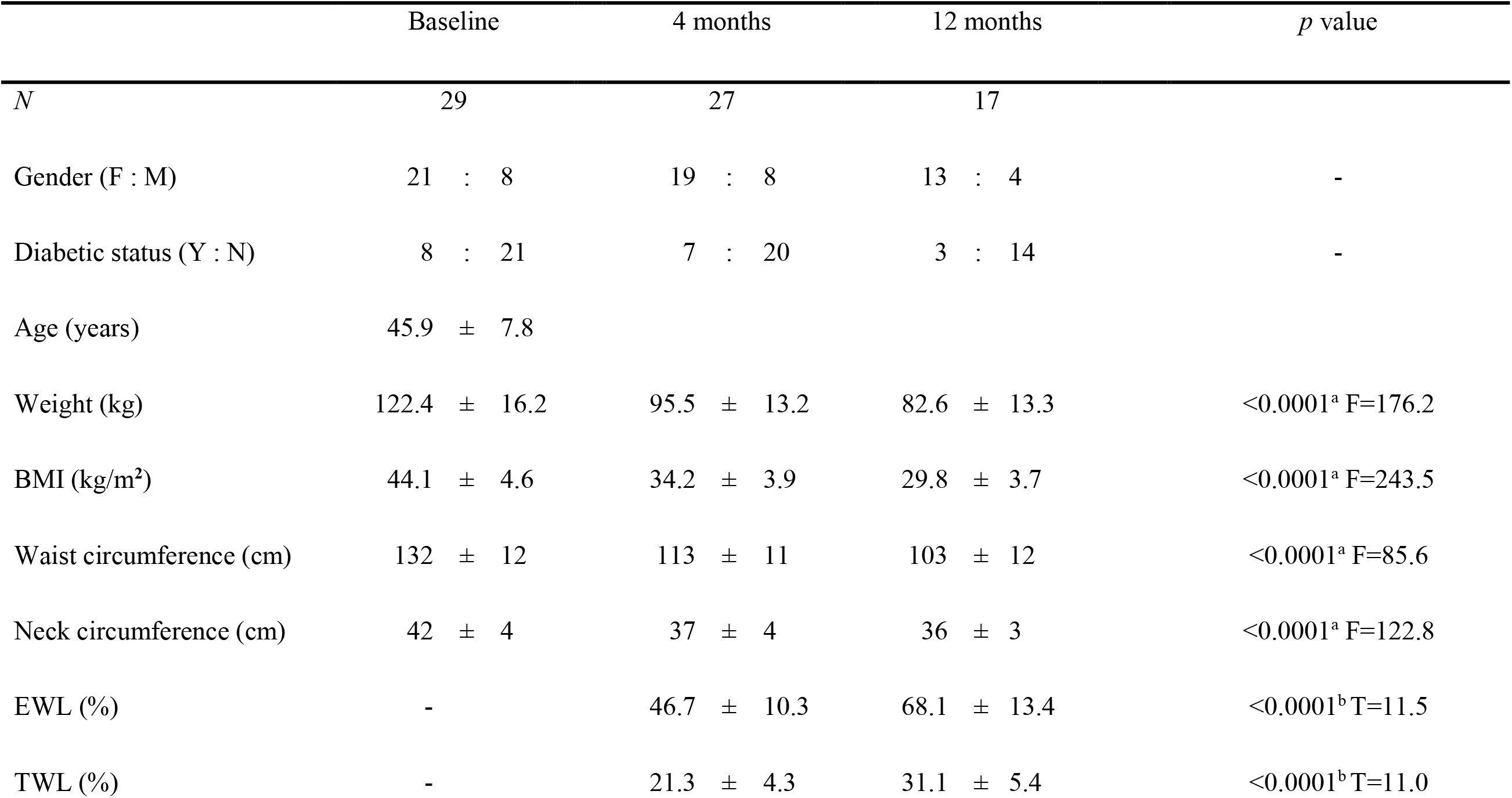
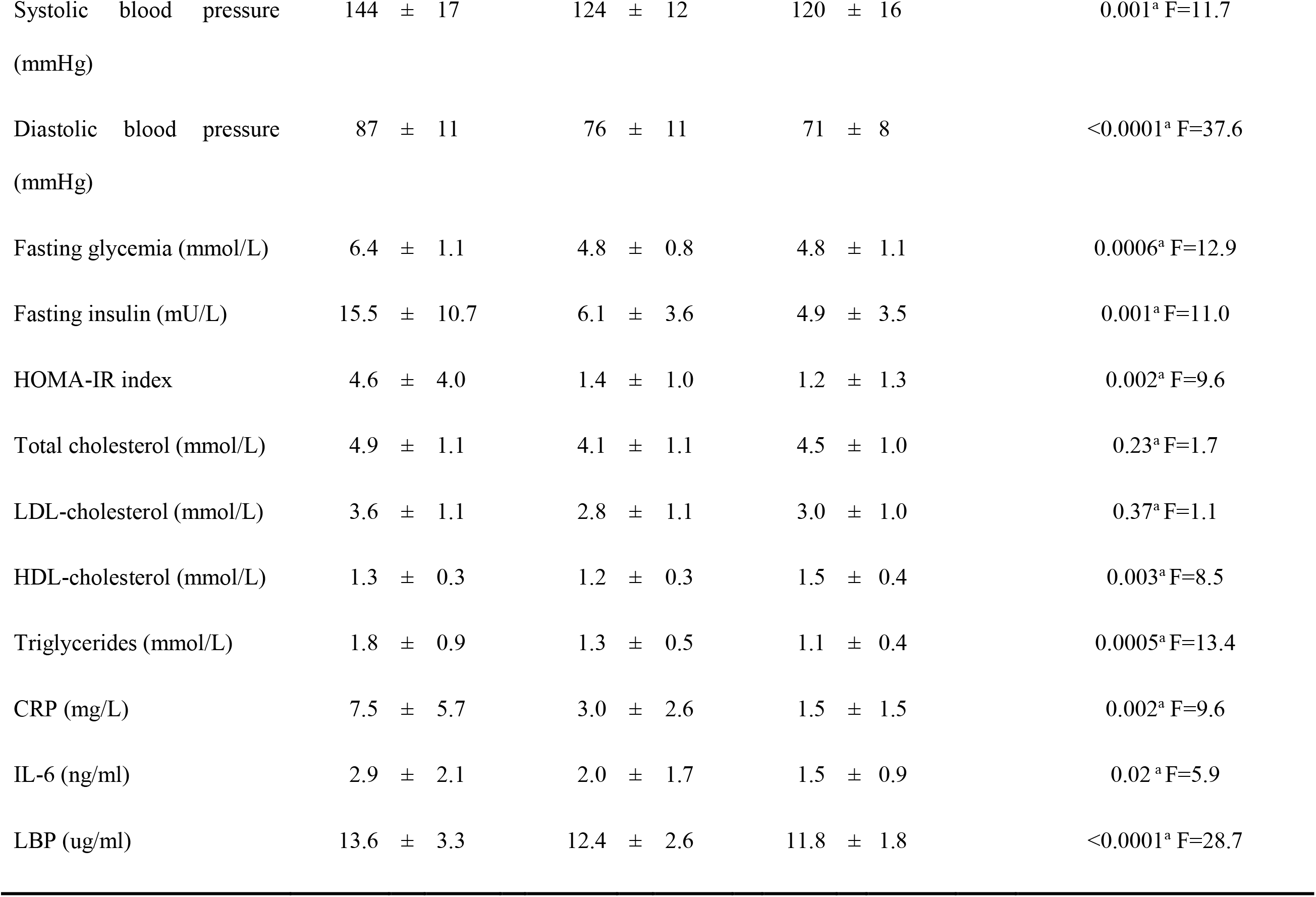
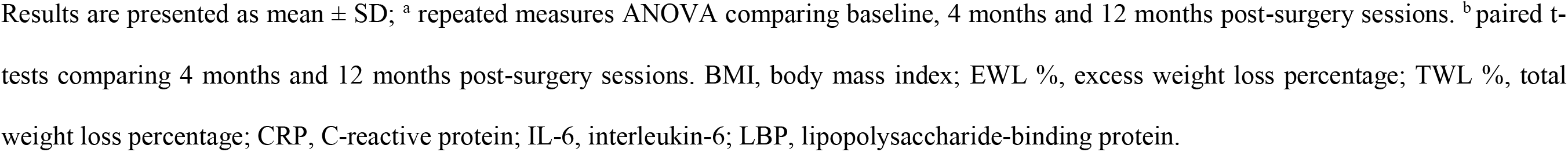
Characteristics of participants at baseline, 4 and 12 months after SG.

|  | Baseline | 4 months | 12 months | *p* value |
| --- | --- | --- | --- | --- |
| *N* | 29 | 27 | 17 |  |
| Gender (F : M) | 21 : 8 | 19 : 8 | 13 : 4 | - |
| Diabetic status (Y : N) | 8 : 21 | 7 : 20 | 3 : 14 | - |
| Age (years) | 45.9 ± 7.8 |  |  |  |
| Weight (kg) | 122.4 ± 16.2 | 95.5 ± 13.2 | 82.6 ± 13.3 | <0.0001<sup>a</sup> F=176.2 |
| BMI (kg/m<sup>2</sup>) | 44.1 ± 4.6 | 34.2 ± 3.9 | 29.8 ± 3.7 | <0.0001<sup>a</sup> F=243.5 |
| Waist circumference (cm) | 132 ± 12 | 113 ± 11 | 103 ± 12 | <0.0001<sup>a</sup> F=85.6 |
| Neck circumference (cm) | 42 ± 4 | 37 ± 4 | 36 ± 3 | <0.0001<sup>a</sup> F=122.8 |
| EWL (%) | - | 46.7 ± 10.3 | 68.1 ± 13.4 | <0.0001<sup>b</sup> T=11.5 |
| TWL (%) | - | 21.3 ± 4.3 | 31.1 ± 5.4 | <0.0001<sup>b</sup> T=11.0 |
| Systolic blood pressure (mmHg) | 144 ± 17 | 124 ± 12 | 120 ± 16 | 0.001<sup>a</sup> F=11.7 |
| Diastolic blood pressure (mmHg) | 87 ± 11 | 76 ± 11 | 71 ± 8 | <0.0001<sup>a</sup> F=37.6 |
| Fasting glycemia (mmol/L) | 6.4 ± 1.1 | 4.8 ± 0.8 | 4.8 ± 1.1 | 0.0006<sup>a</sup> F=12.9 |
| Fasting insulin (mU/L) | 15.5 ± 10.7 | 6.1 ± 3.6 | 4.9 ± 3.5 | 0.001<sup>a</sup> F=11.0 |
| HOMA-IR index | 4.6 ± 4.0 | 1.4 ± 1.0 | 1.2 ± 1.3 | 0.002<sup>a</sup> F=9.6 |
| Total cholesterol (mmol/L) | 4.9 ± 1.1 | 4.1 ± 1.1 | 4.5 ± 1.0 | 0.23<sup>a</sup> F=1.7 |
| LDL-cholesterol (mmol/L) | 3.6 ± 1.1 | 2.8 ± 1.1 | 3.0 ± 1.0 | 0.37<sup>a</sup> F=1.1 |
| HDL-cholesterol (mmol/L) | 1.3 ± 0.3 | 1.2 ± 0.3 | 1.5 ± 0.4 | 0.003<sup>a</sup> F=8.5 |
| Triglycerides (mmol/L) | 1.8 ± 0.9 | 1.3 ± 0.5 | 1.1 ± 0.4 | 0.0005<sup>a</sup> F=13.4 |
| CRP (mg/L) | 7.5 ± 5.7 | 3.0 ± 2.6 | 1.5 ± 1.5 | 0.002<sup>a</sup> F=9.6 |
| IL-6 (ng/ml) | 2.9 ± 2.1 | 2.0 ± 1.7 | 1.5 ± 0.9 | 0.02<sup>a</sup> F=5.9 |
| LBP (ug/ml) | 13.6 ± 3.3 | 12.4 ± 2.6 | 11.8 ± 1.8 | <0.0001<sup>a</sup> F=28.7 |

Results are presented as mean ± SD; <sup>a</sup> repeated measures ANOVA comparing baseline, 4 months and 12 months post-surgery sessions. <sup>b</sup> paired t-tests comparing 4 months and 12 months post-surgery sessions. BMI, body mass index; EWL %, excess weight loss percentage; TWL %, total weight loss percentage; CRP, C-reactive protein; IL-6, interleukin-6; LBP, lipopolysaccharide-binding protein.

### Effect of SG on WM and GM density

**Figure 1** shows the Beta value maps from the voxel-wise mixed effects-models for the WM regions that were significant after FDR correction (p<0.05). A widespread increase in WM density was observed 4 months after SG compared to baseline. These significant increases were found in the cerebellum and cerebellar peduncle but also in the brain stem extending up to cerebral peduncle. There was also a diffuse increase in WM density in the corpus callosum and cingulum projecting out to the corona radiata. Twelve months after the surgery, greater and more significant WM density increases were found compared to baseline in the same significant WM regions observed after the first 4 months, suggesting more pronounced changes after 12 months (**Figure 1**). As expected, significant WM density increases were observed 12 months post-surgery compared to 4 months post-surgery in the same brain regions (p<0.05, after FDR correction, **Figure S1**).

**Figure 1.**
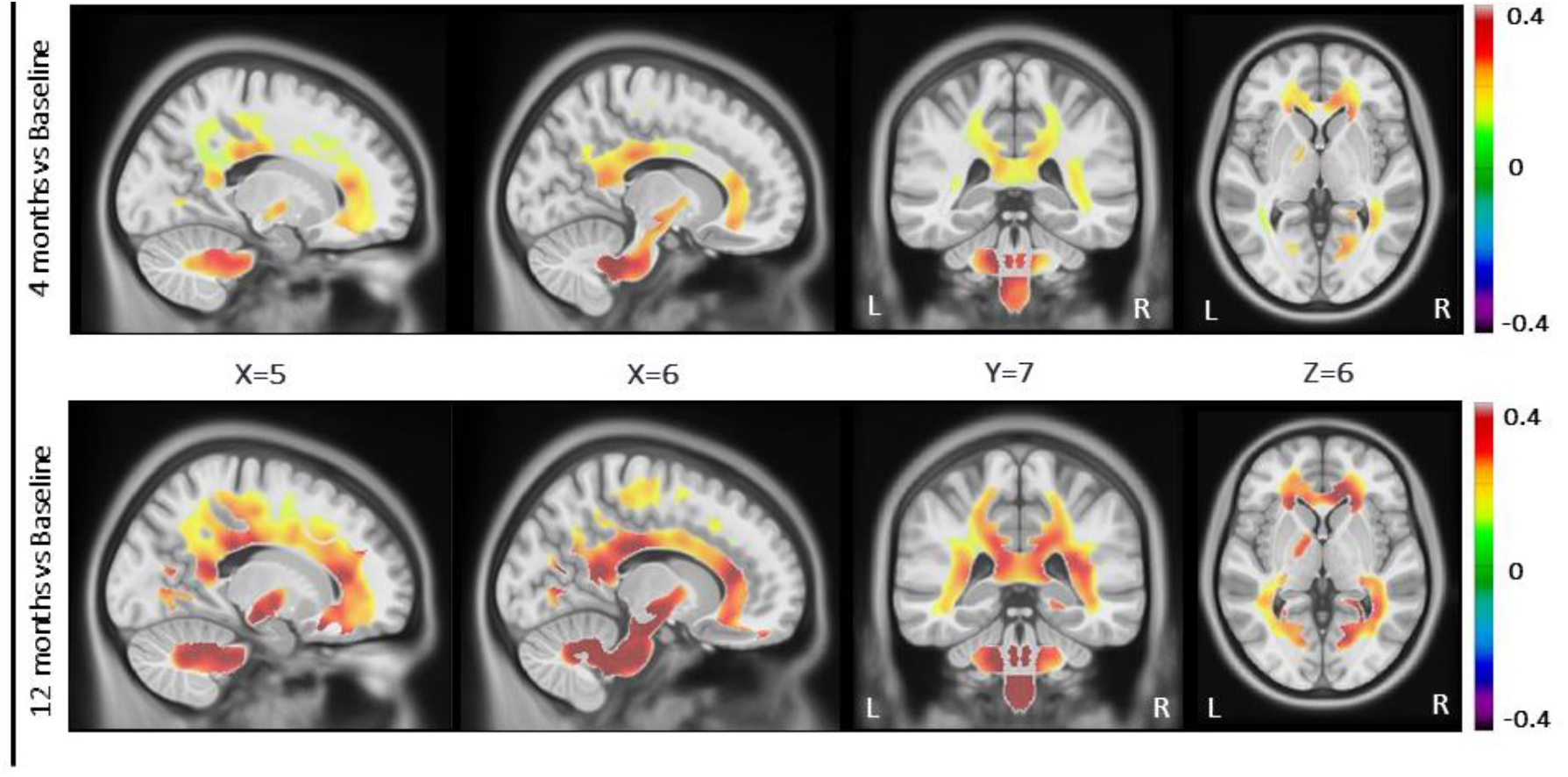
Widespread increase in white matter (WM) density 4 months or 12 months post-surgery compared to baseline. The figure shows the Beta value maps from the voxel-wise mixed effects-models, for the WM regions that were significant after whole brain FDR correction (p<0.05). Warmer colors show higher positive (in red) or negative (in dark purple) Beta values. These results were significant after correcting for age, gender, initial BMI and diabetic status. L, left; R, right

**Figure 2** shows the Beta value maps from the voxel-wise mixed effects-models for the GM regions that were significant after FDR correction (p<0.05). Significant increases in GM density were observed 4 months after SG compared to baseline in several brain regions such as the bilateral occipital cortex, temporal cortex, precentral gyrus and cerebellum as well as right fusiform gyrus, right hippocampus and right insula. Twelve months after SG, greater increases in GM density were found compared to baseline in cortical and subcortical brain regions including bilateral occipital and temporal cortex, precentral gyrus, postcentral gyrus, frontal operculum cortex, fusiform gyrus, insula, parahippocampal gyrus, cerebellum, amygdala, hippocampus and putamen (**Figure 2**). Two regions showed slightly decreased GM density after SG (left cerebellum and right precuneus cortex). No significant difference was observed in GM density comparing the 12-month and the 4-month post-surgery sessions (data not shown).

**Figure 2.**
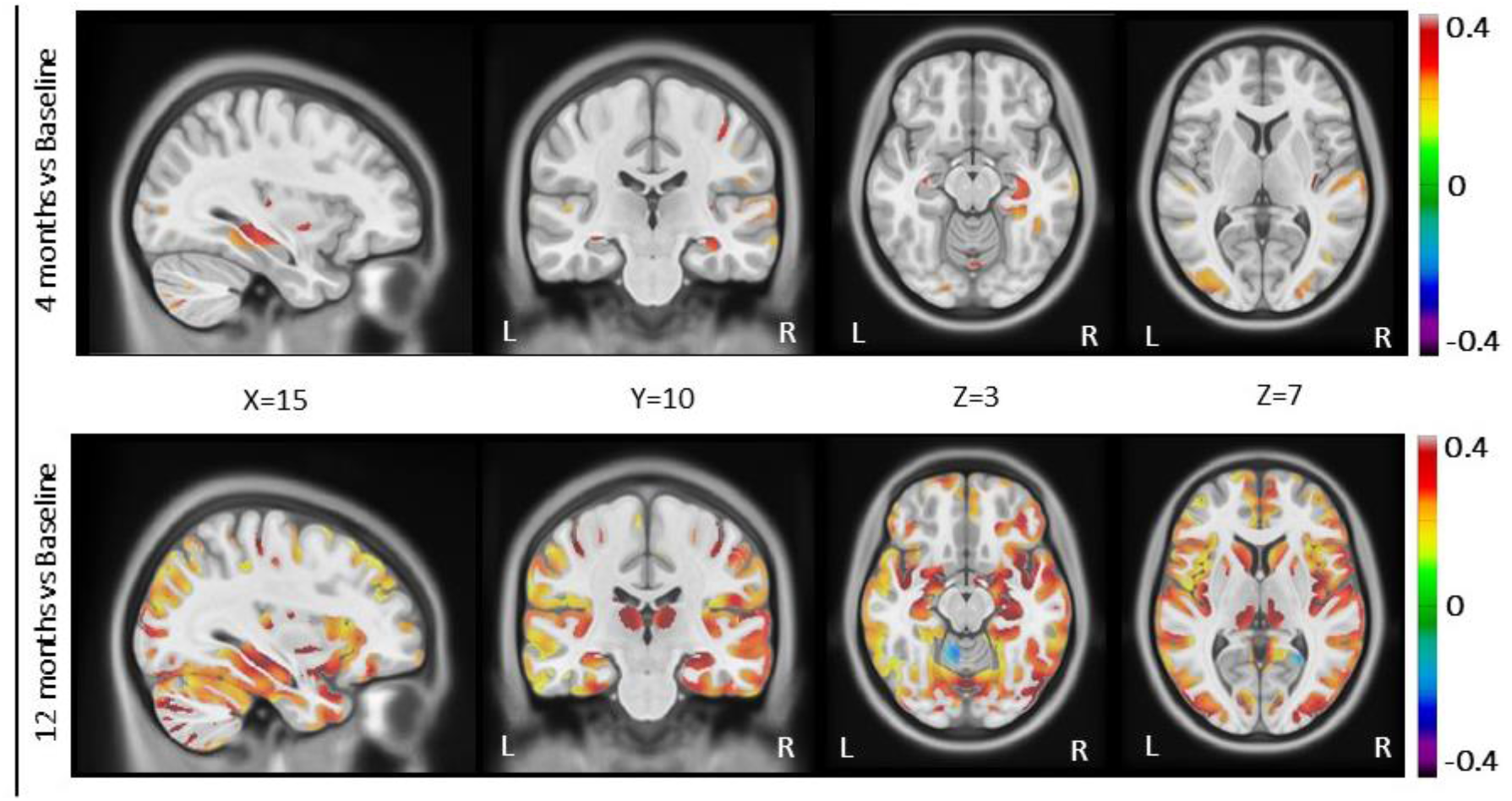
Changes in grey matter (GM) density 4 months or 12 months post-surgery compared to baseline. The figure shows the Beta value maps from the voxel-wise mixed effects-models, for the GM regions that were significant after whole brain FDR correction (p<0.05). Warmer colors show higher positive (in red) or negative (in dark purple) Beta values. These results were significant after correcting for age, gender, initial BMI and diabetic status. L, left; R, right

### Associations between changes in WM density and adiposity, metabolic and inflammatory markers following SG

Mixed-effects models were performed to assess the associations between changes in WM density and adiposity, metabolic or inflammatory variables after SG, accounting for age and gender. Significant associations were found between post-operative reduction in BMI, waist circumference or neck circumference and increased WM density in the cingulum and cerebellar peduncle (p≤0.0001 for all, **Table S1**). A significant association was also observed between post-operative BMI loss and increased WM density in the parietopontine tract (p=0.0016, **Table S1**). Reduced insulin levels following SG were significantly associated with increases in several WM regions such as the acoustic radiation, cingulum, extreme capsule, cerebellar peduncle, occipitopontine tract and parietopontine tract (p≤0.0002 for all, **Table S3**). The same results were observed with the HOMA-IR index. Our linear mixed-effect models showed significant positive associations between HDL-cholesterol levels and WM density in acoustic radiation, extreme capsule, occipitopontine tract, parietopontine tract, anterior commissure, arcuate fasciculus, corpus callosum and corticothalamic pathway (p≤0.0011 for all, **Table S3**). No significant association was found between WM density and circulating levels of glucose, LDL-cholesterol and triglyceride (data not shown). Strong associations were observed between post-operative reductions in LBP and increased WM density in the cingulum, corticospinal tract, extreme capsule, occipitopontine tract, parietopontine tract and spinothalamic tract (p≤0.0017, **Figure 3** and **Table S5**). Decreased plasma IL-6 concentrations were significantly related to increased WM density in cerebellar peduncle and spinothalamic tract only (p≤0.0012, **Table S5**). No significant Bonferroni-corrected associations were found between CRP levels and WM density (**Table S5**).

**Figure 3.**
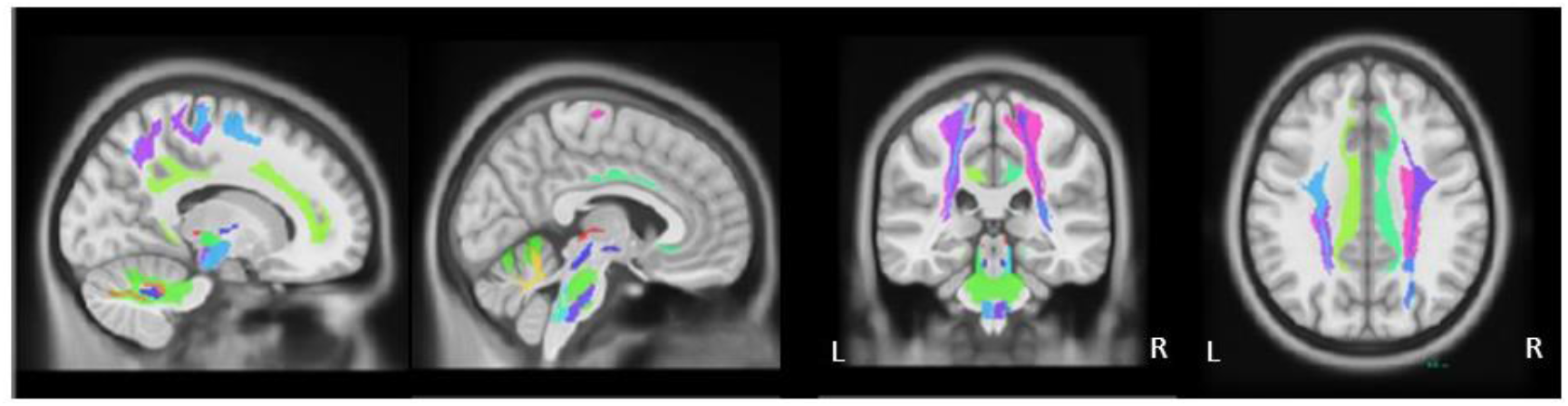
Brain maps of increased white matter regions significantly associated with post-operative reductions in lipopolysaccharide binding protein (LBP) on a standard brain template in Montreal Neurological Institute space. WM regions were extracted from the atlas of Yeh et al. (39). Results of the linear mixed-effect models are presented in Table S5. L, left; R, right

### Associations between changes in GM density and adiposity, metabolic or inflammatory markers following SG

Mixed-effects models were also performed to examine the associations between changes in GM density and adiposity, metabolic or inflammatory variables following SG, accounting for age and gender. Strong associations were observed between post-operative BMI, waist circumference or neck circumference reduction and increased GM density in several cortical and subcortical regions including the precentral gyrus, inferior frontal gyrus, rolandic operculum, insula, hippocampus, amygdala, occipital cortex, fusiform gyrus, angular gyrus, putamen, heschl, temporal cortex, cerebellum and parahipoccampal gyrus (p≤0.0009, **Table S2**). Significant negative associations were found between fasting glucose levels and GM density in bilateral occipital cortex, putamen and cerebellum (p≤0.0006, **Table S4**). Post-operative reductions in insulin levels were significantly associated with increased GM density in the inferior frontal gyrus and rolandic operculum (p≤0.0013, **Table S4**). Post-surgery decreases in triglyceride levels were significantly related to GM density in precentral gyrus, rolandic operculum, supplementary motor area, amygdala, occipital cortex, fusiform gyrus, postcentral gyrus, angular gyrus, temporal cortex and cerebellum (p≤0.0013, **Table S4**). No significant association was observed between changes in GM density and circulating levels of HDL-cholesterol, LDL cholesterol, LBP, IL-6 CRP and HOMA-IR index (data not shown).

### Effect of obesity on WM and GM density: Validation in an independent dataset

To confirm that our results lie in brain areas that are relevant for severe obesity, we used an independent dataset from the HCP. We tested whether the peak of each WM and GM region that significantly increased after the surgery shows BMI-related differences in a separate population. We matched a group of 46 severely obese individuals (BMI: 38.0±2.8 kg/m^2^) with a group of 46 lean individuals (BMI: 22.2±1.3 kg/m^2^) for age, gender and ethnicity (**Table 2**). Mean age was 28.9 years in both groups and most of the individuals were Caucasian (72%). The most significant regions in WM from our clinical sample, such as the brain stem and cerebellar peduncle, had significantly reduced density in obese individuals compared with lean individuals (**Figure 4**, p<0.0001). GM density in the two regions of the cerebellum was significantly reduced in obese individuals relative to controls (**Figure 5**, p<0.0001). To ensure the robustness of the results, mean VBM values were also calculated in a spherical region with each peak as the center and a radius of 2 mm and the analysis was repeated, achieving the same results (data not shown).

**Table 2.** Characteristics of severely obese and normal weight groups.

|  | Severely obese individuals | Normal weight individuals | *p* value |
| --- | --- | --- | --- |
| *N* | 46 | 46 | - |
| Gender (F : M) | 26 : 20 | 26 : 20 | NS |
| Age (years) | 28.9 ± 3.6 | 28.9 ± 3.3 | NS |
| BMI (kg/m<sup>2</sup>) | 38.0 ± 2.8 | 22.2 ± 1.3 | <0.0001 |
| Ethnicity |  |  |  |
| Caucasian (*N*) | 33 | 33 | NS |
| African american (*N*) | 13 | 13 | NS |

Results are presented as mean ± SD; Matched paired t-tests comparing both groups; BMI, body mass index.

**Figure 4.**
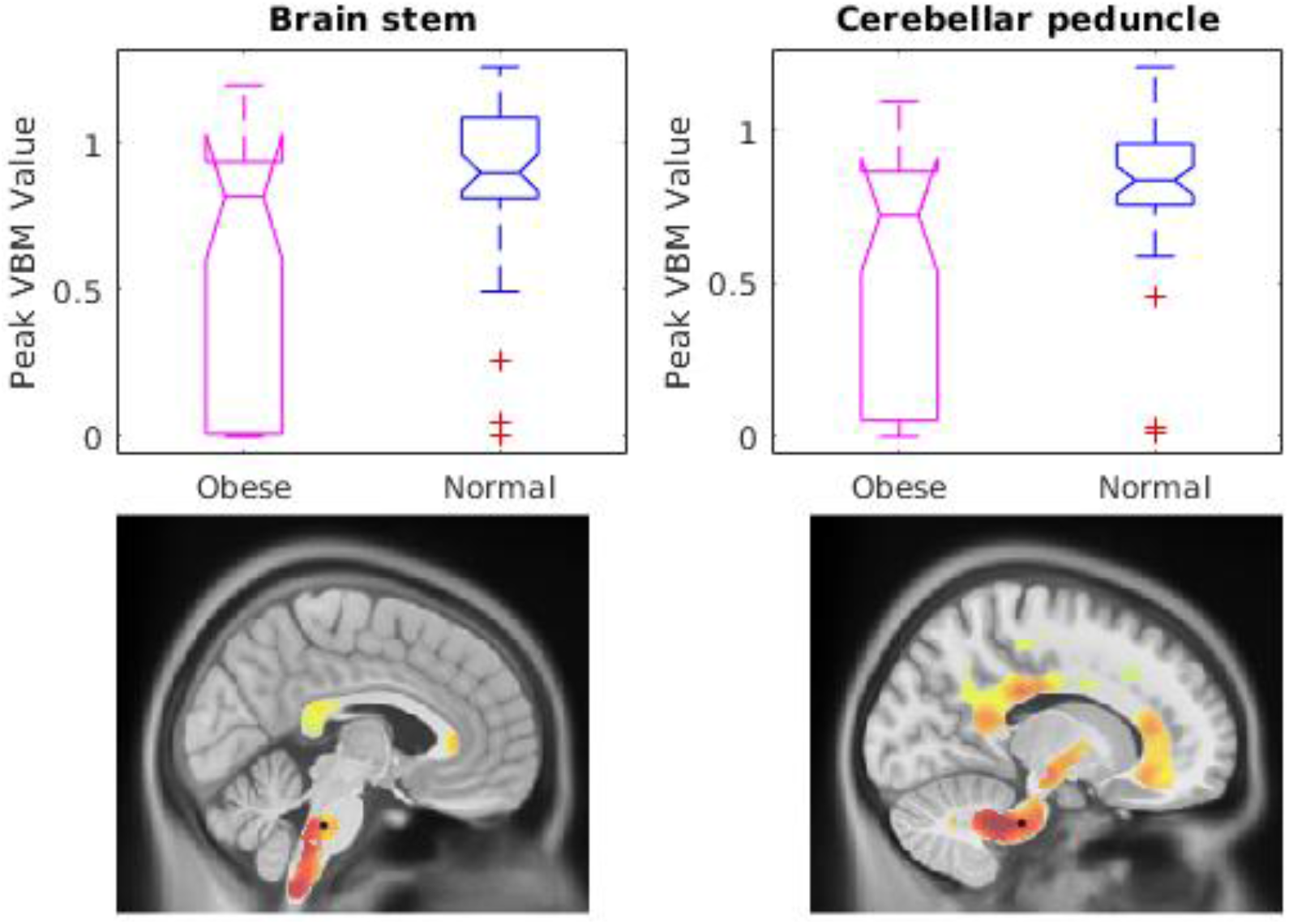
Comparison of the peak Beta value in brain stem and cerebellar peduncle between severely obese individuals and normal weight individuals (p<0.0001). These white matter (WM) regions were significantly increased after bariatric surgery. The localisation of the peak value is identified with a dot on the Beta maps.

**Figure 5.**
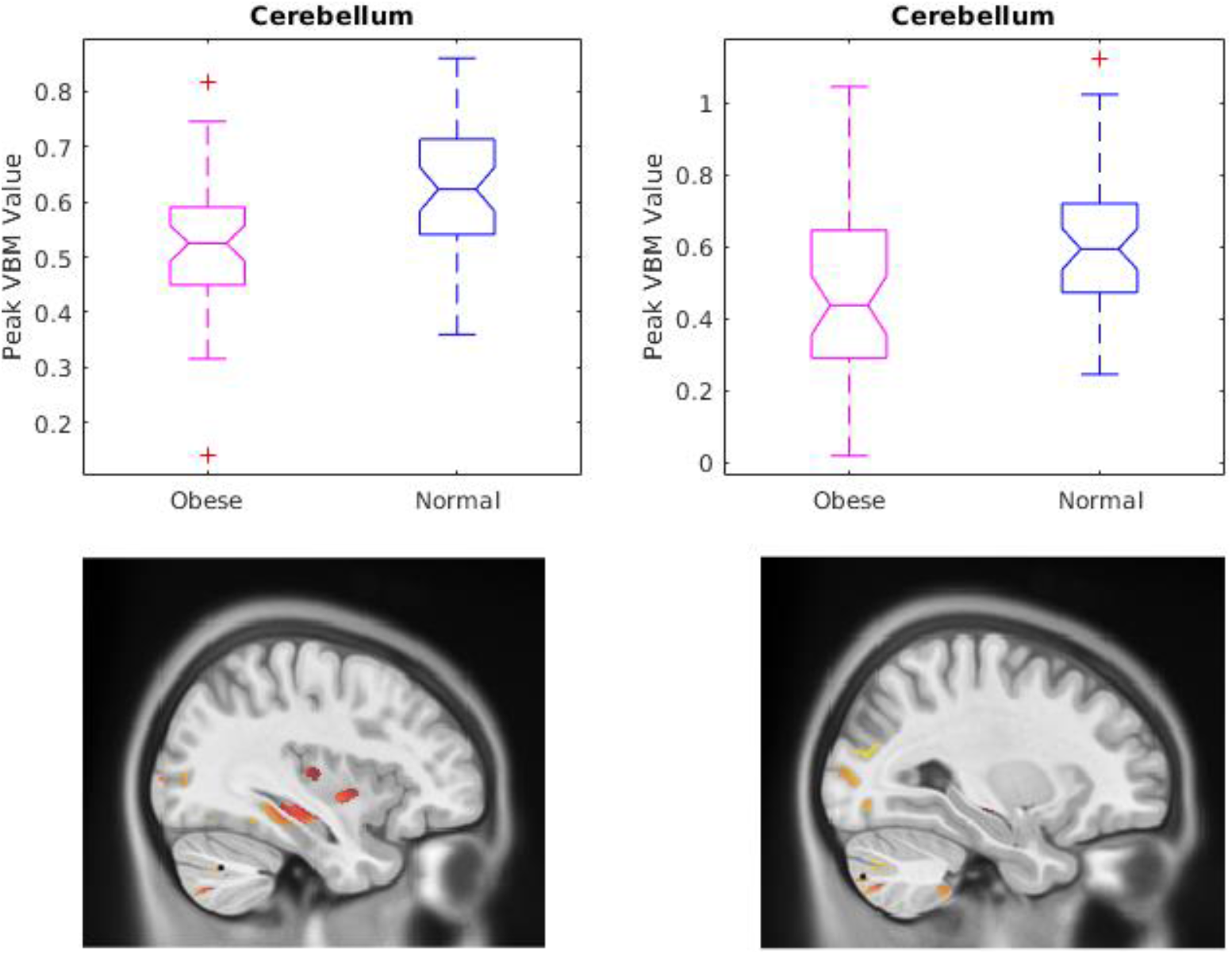
Comparison of the peak Beta value in cerebellum between severely obese individuals and normal weight individuals (p<0.0001). These grey matter (GM) regions were significantly increased after bariatric surgery. The localisation of the peak value is identified with a dot on the Beta value maps.

## DISCUSSION

We tested whether surgery led to recovery in GM and WM density and that these changes are related to improvement in inflammation and metabolic alterations. Overall, we found a widespread increase in WM density and more limited increases in GM density 4 months following SG compared to baseline. These increases were more pronounced and widespread after 12 months and were significantly associated with post-operative weight loss and the metabolic improvements. Our linear mixed-effect models also showed strong associations between post-operative reductions in LBP, a marker of inflammation, and increased WM density. Furthermore, using an independent dataset, we corroborated that these post-operative changes overlapped with baseline brain differences between severely obese and normal-weight participants. More specifically, we found that severe obesity was associated with reduced WM density in the brain stem and cerebellar peduncle as well as reduced GM density in cerebellum, regions that significantly increased after surgery. This last result shows that post-operative brain changes lie in brain areas that are relevant for severe obesity.

As previously reported by other groups (9, 29), we observed that bariatric surgery-induced weight loss was associated with increased global WM density, particularly in the cerebellum, brain stem, cerebellar peduncle, cingulum, corpus callosum and corona radiata. Imaging studies in humans previously reported that obese individuals have reduced WM tissue density and altered WM integrity in dispersed tracts, including the corpus callosum, cingulum, cerebellar peduncle and corona radiata (9, 12, 13, 17, 19, 29, 43). These findings align well with our cross-sectional results showing that WM density in brain stem and cerebellar peduncle, two regions that significantly increased after SG, were reduced in severely obese individuals compared to normal-weight individuals. Interestingly, our results showed that the increases in WM density were more pronounced 12 months post-surgery, suggesting greater WM recovery with the magnitude and duration of weight loss and metabolic improvement. Together, these findings may suggest that weight loss and concomitant improvement of metabolic alterations might reverse obesity-related WM tissue abnormalities.

Such structural increases after surgery were also observed in cortical and subcortical GM regions, mostly in the occipital and temporal/medial temporal lobes (e.g. hippocampus, amygdala, parahippocampal gyrus), precentral gyrus, inferior frontal gyrus, insula, putamen and cerebellum. Some of these changes were detected 4 months following surgery, but they were more extensive after 12 months. As for WM, increases in GM density were significantly associated with the magnitude of weight loss. Our results are consistent with previous bariatric surgery studies on brain morphology showing increased GM density post-surgery, especially in the occipital and temporal regions, fusiform gyrus, postcentral gyrus, inferior frontal gyrus, cerebellum, hippocampus and amygdala (9, 28, 29). Similar observations were found in weight-loss intervention studies (44, 45). Both a recent meta-analysis and studies with large samples demonstrated that obese individuals have reduced GM volumes in several brain regions, including medial prefrontal cortex, inferior frontal gyrus (including insula), cerebellum, temporal lobe (e.g., hippocampal/parahippocampal region and amygdala), precentral gyrus and inferior parietal cortex (8, 46–48). Most of these regions were increased after surgery in our study, which may indirectly suggest a recovery of GM densities in response to bariatric surgery-induced weight loss.

Our findings suggest that changes in WM/GM density after surgery are linked to improvements in cardiometabolic health. More specifically, increases in WM and GM densities were associated with post-operative improvement in glucose homeostasis parameters (fasting insulin and glucose or HOMA-IR index). Furthermore, post-operative increased HDL-cholesterol level was positively associated with increased WM density and post-operative decreased triglyceride level was associated with increased GM density in regions that significantly changed following surgery. These results are consistent with previous studies showing that insulin resistance negatively affects brain function and structure within the hippocampus, dorsolateral prefrontal cortex as well as medial and temporal regions, which may contribute to cognitive impairment (49, 50). There is also some evidence linking dyslipidemia and abnormal lipid metabolism with global and regional GM or WM atrophy or WM hyperintensities (13). Even if the specific contribution of each metabolic change cannot be established in the present study, our results suggest that improvements of glucose homeostasis and dyslipidemia following weight-loss might reverse the obesity-related WM and GM volume alterations.

The chronic, low-grade inflammation related to obesity has also been suggested as a potential mechanism to explain obesity-induced WM alterations (13, 19, 21, 22). More specifically, a recent study has shown that circulating LBP level, a marker of metabolic endotoxemia (51), is associated with decreased integrity in some WM regions and altered cognitive performance in obese participants, suggesting a possible role of LBP in obesity-associated neuroinflammation and neuronal dysfunction (22). In the present study, we found significant reductions in circulating LBP, IL-6 and CRP levels after bariatric surgery, which is consistent with previous studies (52, 53). Moreover, we report strong associations between post-operative reductions in LBP and increased WM density in several regions, including brain stem, cerebellum, cerebral peduncle, cingulum and extreme capsule. Less robust associations were generally observed with reduced post-operative IL-6 levels and no significant association was found with circulating CRP. These results are in line with Rullmann et al. (28) showing no significant association between GM/WM densities or WM diffusivity parameters pre-surgery and changes in CRP post-surgery. Additionally, a recent study reported that circulating CRP and IL-6 levels were not associated with WM integrity parameters or longitudinal change of brain volumes in non-demented elderly participants (54). Even if recent evidence supports the presence of neuroinflammation in the hippocampus, amygdala, cortex and cerebellum in obesity (21), our study revealed no significant association between changes in GM density and postoperative reduction in inflammatory markers. As previously suggested (9), it is possible that WM is more vulnerable to metabolic stress and inflammation associated with obesity. In this vein, Haltia et al. (14) reported that cerebral white matter was more related to visceral fat accumulation, a marker of metabolic dysfunction, rather than total body fat mass. Together, these findings raise the possibility that changes in inflammation and metabolic factors following bariatric surgery improve WM structure and integrity, which may contribute, in part, to the improvement in cognitive functions observed after surgery-induced weight loss (26, 27). A potential mechanism to explain these widespread effects in the brain associated with changes in inflammation and metabolic markers after surgery is the improvement in cerebral blood flow. Studies have consistently shown that bariatric surgery has a cardioprotective effect (55) and improves vascular irrigation in the brain, as shown by decreased carotid intima-media-thickness, a marker of subclinical atherosclerosis, and significant improvement in flow-mediated dilation following surgery (56). Even if the interpretation of the cellular and molecular processes underlying the VBM changes is difficult, we suggest that improved cerebrovascular function and tissue oxygenation following surgery-induced weight loss might lead to axon sprouting, dendritic changes, myelin formation or remodeling, synaptic changes or glial changes, all brain components that may influence MRI signals (57, 58). It is unlikely that increases in GM density are due to neurogenesis since the growth of new neurons is almost inexistent in the human adult brain. Interestingly, Rullman et al. (28) suggested that increased WM density after surgery may result from changes in the composition of fiber tracts instead of structural alteration of axons considering that they found no significant change in WM integrity following bariatric surgery. More studies are needed to examine the changes in WM integrity and brain connectivity following bariatric surgery-induced weight loss.

Some limitations of the study should be acknowledged. The cross-sectional population used to validate our results was younger and did not include participants with metabolic comorbidities, which may have reduced the significant brain difference observed between normal weight and severely obese individuals. Another limitation is that no MR measures of WM integrity (e.g. diffusion imaging) were included in the current study. The fact that all body composition and metabolic and inflammatory measures covary means it is not possible to specify which of these is responsible for recovery of GM and WM density.

In conclusion, bariatric surgery-induced weight loss and improvement in metabolic alterations is associated with widespread increases in WM and GM densities. These post-operative changes overlapped with baseline brain differences between severely obese and normal-weight individuals, which may suggest a recovery of WM and GM alterations after bariatric surgery. Our results also raise the possibility that changes in inflammation or metabolic factors following bariatric surgery improve WM and GM structure. Moreover, in addition to prior work, these findings lend support for bariatric surgery having specific benefits for brain health in morbidly obese individuals.

## ACKNOWLEDGMENTS

We would like to acknowledge the contribution of surgeons, nurses, the medical team of the bariatric surgery program at IUCPQ, MRI technicians, Xavier Moreel, Coordinator of the *Plateforme d’imagerie avancée* at IUCPQ, and Guillaume Gilbert, Engineer, *Phillips* as well as the collaboration of participants. We would like to acknowledge the help of Marie-Frédérique Gauthier for the measurement of cytokines.

## REFERENCES

1. Tchernof A, Despres JP. Pathophysiology of human visceral obesity: an update. Physiol Rev. 2013;93(1):359–404.

2. Despres JP, Lemieux I. Abdominal obesity and metabolic syndrome. Nature. 2006;444(7121):881–7.

3. Albanese E, Launer LJ, Egger M, Prince MJ, Giannakopoulos P, Wolters FJ, et al. Body mass index in midlife and dementia: Systematic review and meta-regression analysis of 589,649 men and women followed in longitudinal studies. Alzheimers Dement (Amst). 2017;8:165–78.

4. Prickett C, Brennan L, Stolwyk R. Examining the relationship between obesity and cognitive function: a systematic literature review. Obes Res Clin Pract. 2015;9(2):93–113.

5. Pedditizi E, Peters R, Beckett N. The risk of overweight/obesity in mid-life and late life for the development of dementia: a systematic review and meta-analysis of longitudinal studies. Age Ageing. 2016;45(1):14–21.

6. Walther K, Birdsill AC, Glisky EL, Ryan L. Structural brain differences and cognitive functioning related to body mass index in older females. Hum Brain Mapp. 2010;31(7):1052–64.

7. Zhang R, Beyer F, Lampe L, Luck T, Riedel-Heller SG, Loeffler M, et al. White matter microstructural variability mediates the relation between obesity and cognition in healthy adults. Neuroimage. 2018;172:239–49.

8. Garcia-Garcia I, Michaud A, Dadar M, Zeighami Y, Neseliler S, Collins DL, et al. Neuroanatomical differences in obesity: meta-analytic findings and their validation in an independent dataset. Int J Obes (Lond). 2018.

9. Tuulari JJ, Karlsson HK, Antikainen O, Hirvonen J, Pham T, Salminen P, et al. Bariatric Surgery Induces White and Grey Matter Density Recovery in the Morbidly Obese: A Voxel-Based Morphometric Study. Hum Brain Mapp. 2016;37(11):3745–56.

10. Ronan L, Alexander-Bloch AF, Wagstyl K, Farooqi S, Brayne C, Tyler LK, et al. Obesity associated with increased brain age from midlife. Neurobiol Aging. 2016;47:63–70.

11. Ho AJ, Raji CA, Becker JT, Lopez OL, Kuller LH, Hua X, et al. Obesity is linked with lower brain volume in 700 AD and MCI patients. Neurobiol Aging. 2010;31(8):1326–39.

12. Kullmann S, Schweizer F, Veit R, Fritsche A, Preissl H. Compromised white matter integrity in obesity. Obes Rev. 2015;16(4):273–81.

13. Alfaro FJ, Gavrieli A, Saade-Lemus P, Lioutas VA, Upadhyay J, Novak V. White matter microstructure and cognitive decline in metabolic syndrome: a review of diffusion tensor imaging. Metabolism. 2018;78:52–68.

14. Haltia LT, Viljanen A, Parkkola R, Kemppainen N, Rinne JO, Nuutila P, et al. Brain white matter expansion in human obesity and the recovering effect of dieting. J Clin Endocrinol Metab. 2007;92(8):3278–84.

15. Raji CA, Ho AJ, Parikshak NN, Becker JT, Lopez OL, Kuller LH, et al. Brain structure and obesity. Hum Brain Mapp. 2010;31(3):353–64.

16. Lampe L, Zhang R, Beyer F, Huhn S, Kharabian-Masouleh S, Preusser S, et al. Visceral Obesity Relates to Deep White Matter Hyperintensities via Inflammation. Ann Neurol. 2018.

17. Kullmann S, Callaghan MF, Heni M, Weiskopf N, Scheffler K, Haring HU, et al. Specific white matter tissue microstructure changes associated with obesity. Neuroimage. 2016;125:36–44.

18. Karlsson HK, Tuulari JJ, Hirvonen J, Lepomaki V, Parkkola R, Hiltunen J, et al. Obesity is associated with white matter atrophy: a combined diffusion tensor imaging and voxel-based morphometric study. Obesity (Silver Spring). 2013;21(12):2530–7.

19. Verstynen TD, Weinstein A, Erickson KI, Sheu LK, Marsland AL, Gianaros PJ. Competing physiological pathways link individual differences in weight and abdominal adiposity to white matter microstructure. Neuroimage. 2013;79:129–37.

20. Thaler JP, Schwartz MW. Minireview: Inflammation and obesity pathogenesis: the hypothalamus heats up. Endocrinology. 2010;151(9):4109–15.

21. Guillemot-Legris O, Muccioli GG. Obesity-Induced Neuroinflammation: Beyond the Hypothalamus. Trends Neurosci. 2017;40(4):237–53.

22. Moreno-Navarrete JM, Blasco G, Puig J, Biarnes C, Rivero M, Gich J, et al. Neuroinflammation in obesity: circulating lipopolysaccharide-binding protein associates with brain structure and cognitive performance. Int J Obes (Lond). 2017;41(11):1627–35.

23. Schauer PR, Kashyap SR, Wolski K, Brethauer SA, Kirwan JP, Pothier CE, et al. Bariatric surgery versus intensive medical therapy in obese patients with diabetes. N Engl J Med. 2012;366(17):1567–76.

24. Gloy VL, Briel M, Bhatt DL, Kashyap SR, Schauer PR, Mingrone G, et al. Bariatric surgery versus non-surgical treatment for obesity: a systematic review and meta-analysis of randomised controlled trials. BMJ. 2013;347:f5934.

25. Sjostrom L, Peltonen M, Jacobson P, Sjostrom CD, Karason K, Wedel H, et al. Bariatric surgery and long-term cardiovascular events. JAMA. 2012;307(1):56–65.

26. Alosco ML, Galioto R, Spitznagel MB, Strain G, Devlin M, Cohen R, et al. Cognitive function after bariatric surgery: evidence for improvement 3 years after surgery. Am J Surg. 2014;207(6):870–6.

27. Handley JD, Williams DM, Caplin S, Stephens JW, Barry J. Changes in Cognitive Function Following Bariatric Surgery: a Systematic Review. Obes Surg. 2016;26(10):2530–7.

28. Rullmann M, Preusser S, Poppitz S, Heba S, Hoyer J, Schutz T, et al. Gastric-bypass surgery induced widespread neural plasticity of the obese human brain. Neuroimage. 2017.

29. Zhang Y, Ji G, Xu M, Cai W, Zhu Q, Qian L, et al. Recovery of brain structural abnormalities in morbidly obese patients after bariatric surgery. Int J Obes (Lond). 2016;40(10):1558–65.

30. Gastrointestinal surgery for severe obesity: National Institutes of Health Consensus Development Conference Statement. Am J Clin Nutr. 1992;55(2 Suppl):615S–9S.

31. Biertho L, Simon-Hould F, Marceau S, Lebel S, Lescelleur O, Biron S. Current Outcomes of Laparoscopic Duodenal Switch. Ann Surg Innov Res. 2016;10:1.

32. Biertho L, Biron S, Hould FS, Lebel S, Marceau S, Marceau P. Is biliopancreatic diversion with duodenal switch indicated for patients with body mass index <50 kg/m2? Surg Obes Relat Dis. 2010;6(5):508–14.

33. Michaud A, Grenier-Larouche T, Caron-Dorval D, Marceau S, Biertho L, Simard S, et al. Biliopancreatic diversion with duodenal switch leads to better postprandial glucose level and beta cell function than sleeve gastrectomy in individuals with type 2 diabetes very early after surgery. Metabolism. 2017;74:10–21.

34. Matthews DR, Hosker JP, Rudenski AS, Naylor BA, Treacher DF, Turner RC. Homeostasis model assessment: insulin resistance and beta-cell function from fasting plasma glucose and insulin concentrations in man. Diabetologia. 1985;28(7):412–9.

35. Michaud A, Drolet R, Noel S, Paris G, Tchernof A. Visceral fat accumulation is an indicator of adipose tissue macrophage infiltration in women. Metabolism. 2012;61(5):689–98.

36. Coupe P, Yger P, Prima S, Hellier P, Kervrann C, Barillot C. An optimized blockwise nonlocal means denoising filter for 3-D magnetic resonance images. IEEE Trans Med Imaging. 2008;27(4):425–41.

37. Sled JG, Zijdenbos AP, Evans AC. A nonparametric method for automatic correction of intensity nonuniformity in MRI data. IEEE Trans Med Imaging. 1998;17(1):87–97.

38. Collins DL, Neelin P, Peters TM, Evans AC. Automatic 3D intersubject registration of MR volumetric data in standardized Talairach space. J Comput Assist Tomogr. 1994;18(2):192–205.

39. Tzourio-Mazoyer N, Landeau B, Papathanassiou D, Crivello F, Etard O, Delcroix N, et al. Automated anatomical labeling of activations in SPM using a macroscopic anatomical parcellation of the MNI MRI single-subject brain. Neuroimage. 2002;15(1):273–89.

40. Yeh FC, Panesar S, Fernandes D, Meola A, Yoshino M, Fernandez-Miranda JC, et al. Population-averaged atlas of the macroscale human structural connectome and its network topology. Neuroimage. 2018;178:57–68.

41. Van Essen DC, Ugurbil K, Auerbach E, Barch D, Behrens TE, Bucholz R, et al. The Human Connectome Project: a data acquisition perspective. Neuroimage. 2012;62(4):2222–31.

42. Glasser MF, Sotiropoulos SN, Wilson JA, Coalson TS, Fischl B, Andersson JL, et al. The minimal preprocessing pipelines for the Human Connectome Project. Neuroimage. 2013;80:105–24.

43. Papageorgiou I, Astrakas LG, Xydis V, Alexiou GA, Bargiotas P, Tzarouchi L, et al. Abnormalities of brain neural circuits related to obesity: A Diffusion Tensor Imaging study. Magn Reson Imaging. 2017;37:116–21.

44. Mueller K, Moller HE, Horstmann A, Busse F, Lepsien J, Bluher M, et al. Physical exercise in overweight to obese individuals induces metabolic- and neurotrophic-related structural brain plasticity. Front Hum Neurosci. 2015;9:372.

45. Prehn K, Jumpertz von Schwartzenberg R, Mai K, Zeitz U, Witte AV, Hampel D, et al. Caloric Restriction in Older Adults-Differential Effects of Weight Loss and Reduced Weight on Brain Structure and Function. Cereb Cortex. 2017;27(3):1765–78.

46. Herrmann MJ, Tesar AK, Beier J, Berg M, Warrings B. Grey matter alterations in obesity: A meta-analysis of whole-brain studies. Obes Rev. 2019;20(3):464–71.

47. Weise CM, Piaggi P, Reinhardt M, Chen K, Savage CR, Krakoff J, et al. The obese brain as a heritable phenotype: a combined morphometry and twin study. Int J Obes (Lond). 2017;41(3):458–66.

48. Vainik U, Baker TE, Dadar M, Zeighami Y, Michaud A, Zhang Y, et al. Neurobehavioral correlates of obesity are largely heritable. Proc Natl Acad Sci U S A. 2018;115(37):9312–7.

49. Heni M, Kullmann S, Preissl H, Fritsche A, Haring HU. Impaired insulin action in the human brain: causes and metabolic consequences. Nat Rev Endocrinol. 2015;11(12):701–11.

50. Cheke LG, Bonnici HM, Clayton NS, Simons JS. Obesity and insulin resistance are associated with reduced activity in core memory regions of the brain. Neuropsychologia. 2017;96:137–49.

51. Laugerette F, Alligier M, Bastard JP, Drai J, Chanseaume E, Lambert-Porcheron S, et al. Overfeeding increases postprandial endotoxemia in men: Inflammatory outcome may depend on LPS transporters LBP and sCD14. Mol Nutr Food Res. 2014;58(7):1513–8.

52. Yang PJ, Lee WJ, Tseng PH, Lee PH, Lin MT, Yang WS. Bariatric surgery decreased the serum level of an endotoxin-associated marker: lipopolysaccharide-binding protein. Surg Obes Relat Dis. 2014;10(6):1182–7.

53. Lindegaard KK, Jorgensen NB, Just R, Heegaard PM, Madsbad S. Effects of Roux-en-Y gastric bypass on fasting and postprandial inflammation-related parameters in obese subjects with normal glucose tolerance and in obese subjects with type 2 diabetes. Diabetol Metab Syndr. 2015;7:12.

54. Gu Y, Vorburger R, Scarmeas N, Luchsinger JA, Manly JJ, Schupf N, et al. Circulating inflammatory biomarkers in relation to brain structural measurements in a non-demented elderly population. Brain Behav Immun. 2017;65:150–60.

55. Gomez-Martin JM, Aracil E, Galindo J, Escobar-Morreale HF, Balsa JA, Botella-Carretero JI. Improvement in cardiovascular risk in women after bariatric surgery as measured by carotid intima-media thickness: comparison of sleeve gastrectomy versus gastric bypass. Surg Obes Relat Dis. 2017;13(5):848–54.

56. Lupoli R, Di Minno MN, Guidone C, Cefalo C, Capaldo B, Riccardi G, et al. Effects of bariatric surgery on markers of subclinical atherosclerosis and endothelial function: a meta-analysis of literature studies. Int J Obes (Lond). 2016;40(3):395–402.

57. Zatorre RJ, Fields RD, Johansen-Berg H. Plasticity in gray and white: neuroimaging changes in brain structure during learning. Nat Neurosci. 2012;15(4):528–36.

58. Wenger E, Brozzoli C, Lindenberger U, Lovden M. Expansion and Renormalization of Human Brain Structure During Skill Acquisition. Trends Cogn Sci. 2017;21(12):930–9.

