## Supplemental data for "Neuroanatomical Changes in White and Grey Matter after Sleeve Gastrectomy"

### Supplementary Materials

**Figure S1.** Increased white matter (WM) density 12 months post-surgery compared to 4 months post-surgery. The figure shows the Beta value maps from the voxel-wise mixed effects-models, for the WM regions that were significant after whole brain FDR correction ($p<0.05$). Warmer colors show higher positive (in red) or negative (in dark purple) Beta values. These results were significant after correcting for age, gender, initial BMI and diabetic status. L, left; R, right

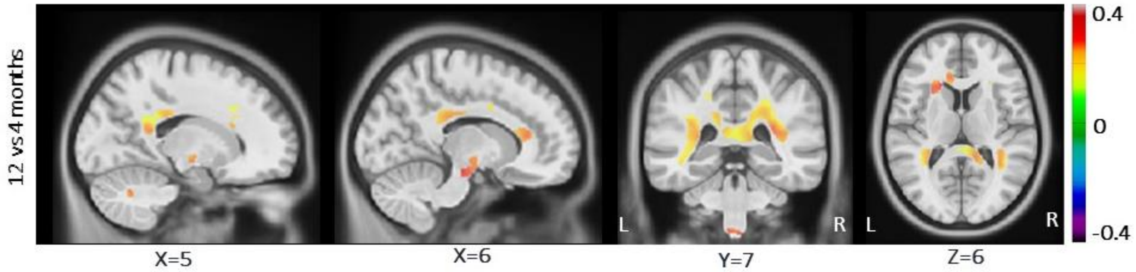

**Table S1.** Summary of the mixed effects models used to assess the association between WM densities and adiposity variables, accounting for age and gender as independent variables.

| Variables | Cingulum | | | Cerebellar peduncle | | | Parietopontine tract | | |
|---|---|---|---|---|---|---|---|---|---|
| | **ß** | **SD** | ***p* value** | **ß** | **SD** | ***p* value** | **ß** | **SD** | ***p* value** |
| Intercept | 0.101 | 0.211 | 0.63 | 0.080 | 0.206 | 0.70 | 0.030 | 0.207 | 0.88 |
| Age | -0.112 | 0.193 | 0.56 | 0.044 | 0.190 | 0.82 | -0.012 | 0.190 | 0.95 |
| Gender | -0.349 | 0.430 | 0.42 | -0.323 | 0.421 | 0.44 | -0.060 | 0.422 | 0.89 |
| BMI | **-0.204** | **0.037** | **<0.0001** | **-0.237** | **0.053** | **<0.0001** | **-0.152** | **0.046** | **0.0016** |
| Intercept | 0.084 | 0.217 | 0.70 | 0.056 | 0.205 | 0.78 | 0.016 | 0.208 | 0.94 |
| Age | -0.082 | 0.198 | 0.68 | 0.055 | 0.188 | 0.77 | 0.007 | 0.191 | 0.97 |
| Gender | -0.305 | 0.444 | 0.49 | -0.248 | 0.418 | 0.56 | -0.024 | 0.426 | 0.954 |
| Waist cir. | **-0.203** | **0.041** | **<0.0001** | **-0.241** | **0.058** | **0.0001** | -0.141 | 0.051 | 0.007 |
| Intercept | -0.006 | 0.221 | 0.98 | -0.068 | 0.194 | 0.73 | -0.049 | 0.211 | 0.82 |
| Age | -0.106 | 0.201 | 0.60 | -0.008 | 0.178 | 0.96 | -0.0189 | 0.194 | 0.92 |
| Gender | 0.090 | 0.464 | 0.846 | 0.290 | 0.416 | 0.49 | 0.263 | 0.448 | 0.56 |
| Neck cir. | **-0.306** | **0.060** | **<0.0001** | **-0.391** | **0.083** | **<0.0001** | -0.216 | 0.074 | 0.005 |

Table shows the regression coefficients (ß), standard deviation (SD) and associated *p* values.
All the variables were z-scored prior to analysis. WM regions were adapted and extracted from Yeh et al. Atlas (40).
waist cir., waist circumference; neck cir., neck circumference; BMI, body mass index.
Significant results after using Bonferroni correction are presented in bold.

**Table S2.** Summary of the mixed effects models used to assess the association between GM densities and adiposity variables, accounting for age and gender as independent variables.

| Variable | Precentral | | | Inferior frontal gyrus, pars opercularis | | | Rolandic operculum | | | Insula | | | Hippocampus | | |
|---|---|---|---|---|---|---|---|---|---|---|---|---|---|---|---|
| | **ß** | **SD** | ***p* value** | **ß** | **SD** | ***p* value** | **ß** | **SD** | ***p* value** | **ß** | **SD** | ***p* value** | **ß** | **SD** | ***p* value** |
| Intercept | 0.206 | 0.206 | 0.32 | 0.324 | 0.186 | 0.08 | 0.135 | 0.201 | 0.505 | 0.148 | 0.188 | 0.43 | 0.353 | 0.197 | 0.08 |
| Age | -0.047 | 0.189 | 0.80 | 0.078 | 0.171 | 0.65 | -0.232 | 0.185 | 0.213 | -0.312 | 0.173 | 0.08 | 0.265 | 0.181 | 0.15 |
| Gender | -0.653 | 0.419 | 0.12 | -1.061 | 0.379 | 0.007 | -0.234 | 0.410 | 0.570 | -0.364 | 0.383 | 0.34 | -1.189 | 0.401 | 0.004 |
| BMI | **-0.270** | **0.071** | **0.0003** | **-0.190** | **0.048** | **0.0002** | **-0.202** | **0.054** | **0.0004** | **-0.211** | **0.060** | **0.0008** | **-0.197** | **0.053** | **0.0005** |
| Intercept | 0.178 | 0.206 | 0.39 | 0.305 | 0.187 | 0.11 | 0.113 | 0.196 | 0.57 | 0.125 | 0.18 | 0.50 | 0.332 | 0.193 | 0.09 |
| Age | -0.040 | 0.190 | 0.83 | 0.088 | 0.172 | 0.61 | -0.231 | 0.180 | 0.20 | -0.309 | 0.169 | 0.07 | 0.269 | 0.178 | 0.13 |
| Gender | -0.559 | 0.420 | 0.19 | -1.001 | 0.382 | 0.01 | -0.160 | 0.400 | 0.69 | -0.287 | 0.376 | 0.45 | -1.119 | 0.395 | 0.006 |
| Waist cir. | **-0.303** | **0.075** | **0.0001** | **-0.196** | **0.052** | **0.0004** | **-0.229** | **0.058** | **0.0002** | **-0.240** | **0.064** | **0.0004** | **-0.216** | **0.057** | **0.0003** |
| Intercept | 0.063 | 0.213 | 0.77 | 0.214 | 0.194 | 0.27 | 0.013 | 0.202 | 0.95 | 0.016 | 0.184 | 0.93 | 0.236 | 0.192 | 0.22 |
| Age | -0.087 | 0.195 | 0.66 | 0.052 | 0.178 | 0.77 | -0.268 | 0.185 | 0.15 | -0.357 | 0.168 | 0.04 | 0.229 | 0.176 | 0.20 |
| Gender | -0.088 | 0.460 | 0.85 | -0.606 | 0.412 | 0.14 | 0.270 | 0.431 | 0.54 | 0.174 | 0.397 | 0.66 | -0.707 | 0.41 | 0.09 |
| Neck cir. | -0.357 | 0.111 | 0.002 | **-0.596** | **0.075** | **0.0002** | **-0.324** | **0.084** | **0.0002** | **-0.343** | **0.09** | **0.0004** | **-0.308** | **0.084** | **0.0005** |

| Variable | Amygdala | | | Occipital cortex | | | Fusiform | | | Angular | | | Putamen | | |
|---|---|---|---|---|---|---|---|---|---|---|---|---|---|---|---|
| | **ß** | **SD** | ***p* value** | **ß** | **SD** | ***p* value** | **ß** | **SD** | ***p* value** | **ß** | **SD** | ***p* value** | **ß** | **SD** | ***p* value** |
| Intercept | 0.304 | 0.217 | 0.16 | 0.347 | 0.180 | 0.06 | 0.135 | 0.196 | 0.49 | 0.273 | 0.195 | 0.16 | 0.160 | 0.171 | 0.35 |
| Age | 0.303 | 0.200 | 0.13 | -0.056 | 0.166 | 0.74 | -0.038 | 0.181 | 0.83 | 0.186 | 0.179 | 0.30 | -0.353 | 0.158 | 0.03 |
| Gender | -0.819 | 0.442 | 0.07 | -1.076 | 0.368 | 0.005 | -0.573 | 0.400 | 0.16 | -1.023 | 0.398 | 0.01 | -0.438 | 0.349 | 0.21 |
| BMI | **-0.223** | **0.054** | **0.0001** | **-0.319** | **0.055** | **<0.0001** | **-0.190** | **0.055** | **0.0009** | **-0.172** | **0.045** | **0.0003** | **-0.291** | **0.070** | **<0.0001** |
| Intercept | 0.279 | 0.212 | 0.19 | 0.312 | 0.175 | 0.08 | 0.115 | 0.195 | 0.56 | 0.256 | 0.195 | 0.19 | 0.130 | 0.169 | 0.44 |
| Age | 0.306 | 0.195 | 0.12 | -0.048 | 0.161 | 0.76 | -0.032 | 0.180 | 0.86 | 0.196 | 0.179 | 0.28 | -0.345 | 0.156 | 0.03 |
| Gender | -0.738 | 0.433 | 0.09 | -0.962 | 0.357 | 0.009 | -0.508 | 0.399 | 0.86 | -0.970 | 0.398 | 0.02 | -0.334 | 0.345 | 0.34 |
| Waist cir. | **-0.248** | **0.058** | **<0.0001** | **-0.354** | **0.058** | **<0.0001** | **-0.203** | **0.059** | **0.0009** | **-0.179** | **0.049** | **0.0005** | **-0.333** | **0.072** | **<0.0001** |
| Intercept | 0.180 | 0.216 | 0.41 | 0.179 | 0.196 | 0.37 | 0.0160 | 0.194 | 0.93 | 0.166 | 0.193 | 0.39 | 0.003 | 0.175 | 0.98 |
| Age | 0.274 | 0.198 | 0.17 | -0.092 | 0.180 | 0.61 | -0.077 | 0.178 | 0.66 | 0.153 | 0.177 | 0.39 | -0.405 | 0.159 | 0.01 |
| Gender | -0.316 | 0.460 | 0.49 | -0.400 | 0.421 | 0.34 | -0.079 | 0.42 | 0.85 | -0.576 | 0.409 | 0.16 | 0.170 | 0.383 | 0.66 |
| Neck cir. | **-0.325** | **0.087** | **0.0004** | **-0.437** | **0.088** | **<0.0001** | **-0.317** | **0.083** | **0.0003** | **-0.289** | **0.070** | **0.0001** | **-0.376** | **0.106** | **0.0007** |

| Variable | Heschl | | | Temporal cortex | | | Cerebellum | | | Parahipoccampal | | |
|---|---|---|---|---|---|---|---|---|---|---|---|---|
| | **ß** | **SD** | ***p* value** | **ß** | **SD** | ***p* value** | **ß** | **SD** | ***p* value** | **ß** | **SD** | ***p* value** |
| Intercept | 0.101 | 0.198 | 0.61 | 0.155 | 0.194 | 0.43 | 0.240 | 0.167 | 0.15 | 0.339 | 0.202 | 0.09 |
| Age | -0.329 | 0.182 | 0.08 | -0.162 | 0.179 | 0.37 | -0.129 | 0.154 | 0.40 | 0.353 | 0.186 | 0.06 |
| Gender | -0.118 | 0.405 | 0.77 | -0.509 | 0.396 | 0.20 | -0.988 | 0.340 | 0.005 | -1.118 | 0.412 | 0.008 |
| BMI | **-0.170** | **0.040** | **<0.0001** | **-0.280** | **0.072** | **0.0002** | **-0.246** | **0.049** | **<0.0001** | -0.188 | 0.058 | 0.0019 |
| Intercept | 0.083 | 0.194 | 0.67 | 0.126 | 0.188 | 0.51 | 0.215 | 0.159 | 0.18 | 0.319 | 0.200 | 0.12 |
| Age | -0.326 | 0.178 | 0.07 | -0.156 | 0.174 | 0.37 | -0.124 | 0.146 | 0.40 | 0.355 | 0.184 | 0.06 |
| Gender | -0.057 | 0.398 | 0.88 | -0.409 | 0.384 | 0.29 | -0.900 | 0.324 | 0.007 | -1.048 | 0.408 | 0.01 |
| Waist cir. | **-0.188** | **0.043** | **<0.0001** | **-0.317** | **0.076** | **<0.0001** | **-0.273** | **0.053** | **<0.0001** | **-0.217** | **0.062** | **0.0008** |
| Intercept | 0.002 | 0.196 | 0.99 | -0.004 | 0.197 | 0.98 | 0.081 | 0.151 | 0.59 | 0.219 | 0.197 | 0.27 |
| Age | -0.351 | 0.180 | 0.06 | -0.2129 | 0.180 | 0.24 | -0.186 | 0.138 | 0.18 | 0.312 | 0.181 | 0.09 |
| Gender | 0.295 | 0.415 | 0.48 | 0.128 | 0.428 | 0.76 | -0.327 | 0.326 | 0.32 | -0.622 | 0.42 | 0.15 |
| Neck cir. | **-0.27** | **0.06** | **<0.0001** | **-0.402** | **0.109** | **<0.0001** | **-0.422** | **0.078** | **<0.0001** | **-0.316** | **0.090** | **0.0007** |

Table shows the regression coefficients (ß), standard deviation (SD) and associated *p* values.
All the variables were z-scored prior to analysis. GM regions adapted and extracted from AAL Atlas (39).
waist cir., waist circumference; neck cir., neck circumference; BMI, body mass index.
Significant results after using Bonferroni correction are presented in bold.

**Table S3.** Summary of the mixed effects models of associations between changes of WM densities and metabolic variables, accounting for age and gender.

| Variable | Acoustic radiation | | | Cingulum | | | Extreme capsule | | | Cerebellar peduncle | | | Occipitopontine tract | | |
|---|---|---|---|---|---|---|---|---|---|---|---|---|---|---|---|
| | **ß** | **SD** | ***p* value** | **ß** | **SD** | ***p* value** | **ß** | **SD** | ***p* value** | **ß** | **SD** | ***p* value** | **ß** | **SD** | ***p* value** |
| Intercept | 0.111 | 0.205 | 0.59 | 0.124 | 0.215 | 0.56 | 0.129 | 0.212 | 0.54 | 0.074 | 0.211 | 0.73 | 0.064 | 0.215 | 0.76 |
| Age | -0.067 | 0.187 | 0.72 | -0.015 | 0.195 | 0.94 | 0.377 | 0.183 | 0.04 | 0.066 | 0.194 | 0.73 | 0.020 | 0.194 | 0.92 |
| Gender | -0.365 | 0.419 | 0.39 | -0.369 | 0.438 | 0.40 | -0.375 | 0.430 | 0.39 | -0.243 | 0.430 | 0.57 | -0.083 | 0.439 | 0.85 |
| Insulin | **-0.194** | **0.050** | **0.0002** | **-0.226** | **0.048** | **0.00001** | **-0.113** | **0.029** | **0.0002** | **-0.264** | **0.067** | **0.0002** | **-0.171** | **0.045** | **0.0004** |
| Intercept | 0.111 | 0.205 | 0.59 | 0.125 | 0.217 | 0.56 | 0.134 | 0.214 | 0.53 | 0.072 | 0.212 | 0.74 | 0.065 | 0.215 | 0.76 |
| Age | -0.044 | 0.187 | 0.81 | 0.015 | 0.196 | 0.94 | 0.411 | 0.183 | 0.03 | 0.087 | 0.195 | 0.66 | 0.045 | 0.194 | 0.82 |
| Gender | -0.372 | 0.419 | 0.38 | -0.382 | 0.443 | 0.39 | -0.399 | 0.433 | 0.36 | -0.246 | 0.433 | 0.57 | -0.094 | 0.439 | 0.83 |
| HOMA-IR | **-0.189** | **0.050** | **0.0004** | **-0.218** | **0.049** | **0.00003** | **-0.107** | **0.029** | **0.0005** | **-0.247** | **0.069** | **0.0007** | **-0.162** | **0.046** | **0.0009** |
| Intercept | 0.075 | 0.210 | 0.72 | 0.091 | 0.214 | 0.67 | 0.128 | 0.214 | 0.55 | 0.036 | 0.219 | 0.87 | 0.042 | 0.22 | 0.85 |
| Age | -0.117 | 0.193 | 0.55 | -0.077 | 0.197 | 0.70 | 0.428 | 0.181 | 0.021 | -0.004 | 0.205 | 0.98 | 0.005 | 0.201 | 0.98 |
| Gender | -0.244 | 0.432 | 0.57 | -0.254 | 0.440 | 0.56 | -0.373 | 0.434 | 0.39 | -0.125 | 0.453 | 0.78 | -0.001 | 0.454 | 1.00 |
| HDL-C | **0.220** | **0.064** | **0.0011** | 0.209 | 0.068 | 0.003 | **0.140** | **0.036** | **0.0002** | 0.210 | 0.090 | 0.02 | **0.201** | **0.057** | **0.0008** |

| | Parietopontine tract | | | Anterior commissure | | | Arcuate fasciculus | | | Corpus callosum | | | Corticothalamic pathway | | |
|---|---|---|---|---|---|---|---|---|---|---|---|---|---|---|---|
| | **ß** | **SD** | ***p* value** | **ß** | **SD** | ***p* value** | **ß** | **SD** | ***p* value** | **ß** | **SD** | ***p* value** | **ß** | **SD** | ***p* value** |
| Intercept | 0.034 | 0.205 | 0.87 | -0.007 | 0.199 | 0.97 | 0.075 | 0.216 | 0.72 | 0.052 | 0.208 | 0.80 | 0.082 | 0.207 | 0.69 |
| Age | 0.10 | 0.187 | 0.96 | -0.146 | 0.184 | 0.43 | -0.04 | 0.185 | 0.82 | -0.101 | 0.190 | 0.60 | -0.108 | 0.189 | 0.57 |
| Gender | -0.006 | 0.418 | 0.99 | -0.07 | 0.406 | 0.85 | -0.207 | 0.438 | 0.64 | -0.0831 | 0.425 | 0.85 | -0.191 | 0.421 | 0.65 |
| Insulin | **-0.204** | **0.055** | **0.0004** | -0.162 | 0.069 | 0.021 | -0.08 | 0.028 | 0.006 | -0.160 | 0.052 | 0.003 | -0.133 | 0.057 | 0.02 |
| Intercept | 0.034 | 0.205 | 0.87 | -0.009 | 0.201 | 0.96 | 0.078 | 0.216 | 0.72 | 0.052 | 0.209 | 0.80 | 0.082 | 0.207 | 0.69 |
| Age | 0.031 | 0.187 | 0.87 | -0.134 | 0.185 | 0.47 | -0.021 | 0.184 | 0.91 | -0.082 | 0.190 | 0.67 | -0.094 | 0.189 | 0.62 |
| Gender | -0.011 | 0.418 | 0.98 | -0.074 | 0.410 | 0.86 | -0.221 | 0.438 | 0.62 | -0.089 | 0.426 | 0.84 | -0.193 | 0.422 | 0.65 |
| HOMA-IR | **-0.198** | **0.056** | **0.0007** | -0.161 | 0.069 | 0.02 | -0.081 | 0.028 | 0.006 | -0.158 | 0.053 | 0.004 | -0.133 | 0.057 | 0.02 |
| Intercept | -0.003 | 0.204 | 0.99 | -0.063 | 0.197 | 0.75 | 0.065 | 0.214 | 0.76 | 0.020 | 0.207 | 0.92 | 0.047 | 0.204 | 0.82 |
| Age | -0.060 | 0.189 | 0.75 | -0.270 | 0.184 | 0.15 | -0.332 | 0.177 | 0.85 | -0.160 | 0.191 | 0.40 | -0.187 | 0.189 | 0.32 |
| Gender | 0.140 | 0.420 | 0.74 | 0.151 | 0.407 | 0.71 | -0.161 | 0.433 | 0.71 | 0.0494 | 0.426 | 0.91 | -0.031 | 0.420 | 0.94 |
| HDL-C | **0.245** | **0.070** | **0.0008** | **0.286** | **0.080** | **0.0007** | **0.140** | **0.032** | **0.00005** | **0.221** | **0.064** | **0.0011** | **0.229** | **0.067** | **0.0012** |

Table shows the regression coefficients (ß), standard deviation (SD) and associated *p* values.
All the variables were z-scored prior to analysis. WM regions were adapted and extracted from Yeh et al. Atlas (40).
Significant results after using Bonferroni correction are presented in bold.

**Table S4**. Summary of the mixed effects models of associations between changes of GM densities and metabolic variables, accounting for age and gender.

| Variable | Precentral | | | Inferior frontal gyrus, pars opercularis | | | Rolandic operculum | | | Supplementary motor area | | | Amygdala | | |
|---|---|---|---|---|---|---|---|---|---|---|---|---|---|---|---|
| | ß | SD | *p* value | ß | SD | *p* value | ß | SD | *p* value | ß | SD | *p* value | ß | SD | *p* value |
| Intercept | 0.216 | 0.206 | 0.30 | 0.270 | 0.192 | 0.16 | 0.121 | 0.203 | 0.56 | 0.153 | 0.201 | 0.45 | 0.296 | 0.213 | 0.17 |
| Age | 0.034 | 0.191 | 0.86 | 0.131 | 0.176 | 0.46 | -0.168 | 0.188 | 0.37 | -0.146 | 0.185 | 0.44 | 0.368 | 0.196 | 0.07 |
| Gender | -0.627 | 0.419 | 0.14 | -1.039 | 0.392 | 0.01 | -0.236 | 0.416 | 0.57 | -0.523 | 0.411 | 0.21 | -0.785 | 0.436 | 0.08 |
| Glucose | -0.289 | 0.087 | 0.0015 | -0.146 | 0.056 | 0.01 | -0.203 | 0.067 | 0.003 | -0.137 | 0.068 | 0.05 | -0.228 | 0.068 | 0.0015 |
| Intercept | 0.211 | 0.210 | 0.32 | 0.272 | 0.188 | 0.15 | 0.123 | 0.194 | 0.53 | 0.156 | 0.202 | 0.44 | 0.286 | 0.215 | 0.19 |
| Age | -0.026 | 0.195 | 0.89 | 0.102 | 0.173 | 0.56 | -0.207 | 0.179 | 0.25 | -0.166 | 0.187 | 0.38 | 0.320 | 0.198 | 0.11 |
| Gender | -0.648 | 0.427 | 0.13 | -1.034 | 0.383 | 0.01 | -0.239 | 0.395 | 0.55 | -0.526 | 0.412 | 0.21 | -0.793 | 0.438 | 0.08 |
| Insulin | -0.181 | 0.096 | 0.06 | **-0.192** | **0.056** | **0.0012** | **-0.236** | **0.070** | **0.0013** | -0.176 | 0.069 | 0.01 | -0.158 | 0.076 | 0.04 |
| Intercept | 0.239 | 0.209 | 0.26 | 0.283 | 0.191 | 0.14 | 0.137 | 0.205 | 0.51 | 0.164 | 0.189 | 0.39 | 0.305 | 0.204 | 0.14 |
| Age | 0.064 | 0.194 | 0.74 | 0.150 | 0.175 | 0.39 | -0.145 | 0.189 | 0.44 | -0.128 | 0.174 | 0.47 | 0.372 | 0.188 | 0.05 |
| Gender | -0.064 | 0.194 | 0.74 | -1.040 | 0.391 | 0.01 | -0.246 | 0.418 | 0.56 | -0.502 | 0.386 | 0.20 | -0.778 | 0.415 | 0.06 |
| TG | **-0.390** | **0.096** | **0.00015** | -0.217 | 0.065 | 0.0015 | **-0.270** | **0.078** | **0.0010** | **-0.264** | **0.078** | **0.0012** | **-0.29** | **0.083** | **0.0009** |

| Variable | Occipital cortex | | | Fusiform | | | Postcentral | | | Angular | | | Putamen | | |
|---|---|---|---|---|---|---|---|---|---|---|---|---|---|---|---|
| | ß | SD | *p* value | ß | SD | *p* value | ß | SD | *p* value | ß | SD | *p* value | ß | SD | *p* value |
| Intercept | 0.296 | 0.191 | 0.13 | 0.113 | 0.202 | 0.58 | 0.224 | 0.191 | 0.24 | 0.226 | 0.194 | 0.25 | 0.111 | 0.169 | 0.51 |
| Age | 0.007 | 0.177 | 0.97 | 0.028 | 0.186 | 0.88 | 0.056 | 0.180 | 0.75 | 0.248 | 0.178 | 0.17 | -0.321 | 0.157 | 0.04 |
| Gender | -0.992 | 0.391 | 0.01 | -0.569 | 0.413 | 0.17 | -0.676 | 0.388 | 0.09 | -0.979 | 0.396 | 0.02 | -0.307 | 0.343 | 0.37 |
| Glucose | **-0.286** | **0.073** | **0.0002** | -0.180 | 0.067 | 0.009 | -0.238 | 0.083 | 0.006 | -0.162 | 0.054 | 0.004 | **-0.318** | **0.084** | **0.0003** |
| Intercept | 0.289 | 0.192 | 0.14 | 0.109 | 0.192 | 0.57 | 0.220 | 0.197 | 0.27 | 0.227 | 0.197 | 0.25 | 0.100 | 0.175 | 0.57 |
| Age | -0.054 | 0.178 | 0.76 | -0.016 | 0.177 | 0.93 | 0.008 | 0.182 | 0.97 | 0.227 | 0.181 | 0.21 | -0.394 | 0.162 | 0.02 |
| Gender | -1.005 | 0.391 | 0.01 | -0.555 | 0.392 | 0.16 | -0.699 | 0.399 | 0.09 | -0.999 | 0.402 | 0.02 | -0.322 | 0.354 | 0.37 |
| Insulin | -0.219 | 0.082 | 0.009 | -0.216 | 0.069 | 0.003 | -0.149 | 0.090 | 0.10 | -0.134 | 0.059 | 0.03 | -0.187 | 0.093 | 0.05 |
| Intercept | 0.310 | 0.190 | 0.11 | 0.127 | 0.193 | 0.51 | 0.244 | 0.212 | 0.25 | 0.236 | 0.186 | 0.21 | 0.115 | 0.165 | 0.49 |
| Age | 0.013 | 0.176 | 0.94 | 0.044 | 0.178 | 0.80 | 0.102 | 0.195 | 0.60 | 0.257 | 0.170 | 0.14 | -0.341 | 0.154 | 0.03 |
| Gender | -1.005 | 0.388 | 0.12 | -0.560 | 0.393 | 0.16 | -0.379 | 0.431 | 0.12 | -0.974 | 0.379 | 0.012 | -0.316 | 0.335 | 0.35 |
| TG | **-0.322** | **0.087** | **0.0005** | **-0.282** | **0.077** | **0.0005** | **-0.377** | **0.088** | **0.00007** | **-0.230** | **0.065** | **0.0008** | -0.294 | 0.097 | 0.004 |

| Variable | Temporal cortex | | | Cerebellum | | |
|---|---|---|---|---|---|---|
| | **ß** | **SD** | ***p* value** | **ß** | **SD** | ***p* value** |
| Intercept | 0.136 | 0.193 | 0.49 | 0.194 | 0.168 | 0.25 |
| Age | -0.112 | 0.179 | 0.54 | -0.085 | 0.156 | 0.59 |
| Gender | -0.459 | 0.393 | 0.25 | -0.931 | 0.343 | 0.01 |
| Glucose | -0.270 | 0.091 | 0.004 | **-0.228** | **0.063** | **0.0006** |
| Intercept | 0.145 | 0.192 | 0.45 | 0.193 | 0.171 | 0.26 |
| Age | -0.146 | 0.178 | 0.41 | -0.126 | 0.158 | 0.44 |
| Gender | -0.486 | 0.389 | 0.22 | -0.942 | 0.348 | 0.009 |
| Insulin | -0.287 | 0.094 | 0.003 | -0.202 | 0.067 | 0.004 |
| Intercept | 0.149 | 0.186 | 0.42 | 0.207 | 0.171 | 0.23 |
| Age | -0.106 | 0.173 | 0.54 | -0.074 | 0.158 | 0.64 |
| Gender | -0.462 | 0.378 | 0.22 | -0.948 | 0.349 | 0.008 |
| TG | **-0.344** | **0.101** | **0.0012** | **-0.252** | **0.075** | **0.0013** |

The table shows the regression coefficients (ß), standard deviation (SD) and associated *p* values.
All the variables were z-scored prior to analysis. GM regions adapted and extracted from AAL Atlas (39).
TG, triglycerides. Significant results after using Bonferroni correction are presented in bold.

**Table S5**. Summary of the mixed-effects models used to assess the associations between WM densities and plasma inflammatory variables, accounting for age and gender as independent variables.

| Variable | Cingulum | | | Corticospinal tract | | | Extreme Capsule | | | Cerebellar peduncle | | | Occipitopontine tract | | |
|---|---|---|---|---|---|---|---|---|---|---|---|---|---|---|---|
| | ß | SD | *p* value | ß | SD | *p* value | ß | SD | *p* value | ß | SD | *p* value | ß | SD | *p* value |
| Intercept | 0.074 | 0.220 | 0.74 | 0.066 | 0.209 | 0.75 | 0.104 | 0.217 | 0.63 | 0.004 | 0.213 | 0.99 | 0.015 | 0.213 | 0.95 |
| Age | -0.080 | 0.203 | 0.69 | -0.094 | 0.194 | 0.63 | 0.366 | 0.189 | 0.06 | -0.04 | 0.198 | 0.84 | -0.046 | 0.196 | 0.82 |
| Gender | -0.381 | 0.447 | 0.40 | -0.303 | 0.426 | 0.48 | -0.398 | 0.438 | 0.37 | -0.24 | 0.434 | 0.58 | -0.076 | 0.434 | 0.86 |
| LBP | **-0.181** | **0.053** | **0.0010** | **-0.224** | **0.060** | **0.0004** | **-0.100** | **0.030** | **0.0015** | **-0.279** | **0.067** | **<0.0001** | **-0.164** | **0.047** | **0.0009** |
| Intercept | 0.154 | 0.217 | 0.48 | 0.139 | 0.206 | 0.50 | 0.151 | 0.215 | 0.48 | 0.101 | 0.213 | 0.637 | 0.079 | 0.219 | 0.72 |
| Age | -0.019 | 0.198 | 0.92 | -0.005 | 0.189 | 0.98 | 0.427 | 0.188 | 0.03 | 0.055 | 0.196 | 0.78 | 0.053 | 0.199 | 0.79 |
| Gender | 0.500 | 0.441 | 0.26 | -0.402 | 0.420 | 0.34 | -0.480 | 0.436 | 0.28 | -0.377 | 0.435 | 0.39 | -0.185 | 0.446 | 0.68 |
| CRP | -0.137 | 0.056 | 0.02 | -0.080 | 0.069 | 0.25 | -0.061 | 0.033 | 0.07 | -0.146 | 0.076 | 0.06 | -0.068 | 0.052 | 0.19 |
| Intercept | 0.439 | 0.256 | 0.09 | 0.449 | 0.248 | 0.08 | 0.275 | 0.231 | 0.24 | 0.532 | 0.255 | 0.04 | 0.345 | 0.244 | 0.16 |
| Age | 0.004 | 0.227 | 0.99 | 0.038 | 0.213 | 0.86 | 0.462 | 0.202 | 0.03 | 0.133 | 0.216 | 0.54 | 0.061 | 0.217 | 0.78 |
| Gender | -0.423 | 0.477 | 0.38 | -0.347 | 0.441 | 0.44 | -0.427 | -0.449 | 0.35 | -0.296 | 0.446 | 0.51 | -0.118 | 0.459 | 0.80 |
| IL-6 | -0.130 | 0.044 | 0.004 | -0.146 | 0.051 | 0.006 | -0.071 | -0.026 | 0.008 | **-0.205** | **0.055** | **0.0004** | -0.125 | 0.040 | 0.003 |

| Variable | Parietopontine Tract | | | Spinothalamic tract | | |
|---|---|---|---|---|---|---|
| | ß | SD | *p* value | ß | SD | *p* value |
| Intercept | 0.019 | 0.213 | 0.93 | 0.046 | 0.184 | 0.80 |
| Age | -0.057 | 0.198 | 0.77 | -0.248 | 0.172 | 0.16 |
| Gender | -0.013 | 0.435 | 0.98 | -0.199 | 0.374 | 0.60 |
| LBP | **-0.200** | **0.054** | **0.0005** | **-0.312** | **0.095** | **0.0017** |
| Intercept | 0.051 | 0.209 | 0.81 | 0.097 | 0.189 | 0.61 |
| Age | 0.021 | 0.191 | 0.91 | -0.167 | 0.173 | 0.34 |
| Gender | -0.109 | 0.426 | 0.80 | -0.236 | 0.387 | 0.54 |
| CRP | -0.085 | 0.062 | 0.17 | -0.056 | 0.105 | 0.59 |
| Intercept | 0.310 | 0.243 | 0.21 | 0.631 | 0.246 | 0.01 |
| Age | 0.077 | 0.211 | 0.72 | -0.103 | 0.186 | 0.58 |
| Gender | -0.052 | 0.440 | 0.91 | -0.227 | 0.378 | 0.55 |
| IL-6 | -0.129 | 0.047 | 0.008 | **-0.229** | **0.067** | **0.0012** |

The table shows the regression coefficients (ß), standard deviation (SD) and associated *p* values.
All the variables were z-scored prior to analysis. WM regions were adapted and extracted from Yeh et al. Atlas (40).
Significant results after using Bonferroni correction are presented in bold.
